# Deorphanisation of novel biogenic amine-gated ion channels identifies a new serotonin receptor for learning

**DOI:** 10.1101/2020.09.17.301382

**Authors:** Julia Morud, Iris Hardege, He Liu, Taihong Wu, Swaraj Basu, Yun Zhang, William R Schafer

## Abstract

Pentameric ligand-gated ion channels (LGCs) play conserved, critical roles in fast synaptic transmission, and changes in LGC expression and localisation are thought to underlie many forms of learning and memory. The *C. elegans* genome encodes a large number of LGCs without a known ligand or function. Here, we deorphanize five members of a family of Cys-loop LGCs by characterizing their diverse functional properties that are activated by biogenic amine neurotransmitters. To analyse the neuronal function of these LGCs, we show that a novel serotonin-gated cation channel, LGC-50, is essential for aversive olfactory learning. *lgc-50* mutants show a specific defect in learned olfactory avoidance of pathogenic bacteria, a process known to depend on serotonergic neurotransmission. Remarkably, the expression of LGC-50 in neuronal processes is enhanced by olfactory conditioning; thus, the regulated expression of these receptors at synapses appears to represent a molecular cornerstone of the learning mechanism.

## Introduction

Synaptic plasticity, the selective strengthening or weakening of individual synaptic connections, is fundamental to the diverse forms of learning and memory in all animals. At the molecular and cellular levels, most forms of synaptic plasticity are thought to involve alterations in the abundance, density or sensitivity of ionotropic neurotransmitter receptors at the postsynaptic membrane. These receptors fall into two general types: the tetrameric glutamate (GluR) receptors, which mediate much excitatory transmission in the vertebrate brain, and the pentameric Cys-loop receptors, which include anion-selective GABA_A_, glycine receptors and cation-selective nicotinic acetylcholine and 5-HT_3_ serotonin receptors. Although much research has emphasised the roles of GluRs in synaptic plasticity mechanisms, increasing evidence suggests that the regulation of Cys-lop channels is also important. For example, regulation of 5-HT_3_ receptor expression and abundance has been implicated in a variety of serotonin-dependent learning processes, such as reward, fear extinction, and cross-modal plasticity following sensory loss (Kondo et al., 2014; Wang et al., 2019; Zhang et al., 2005). However, the molecular mechanisms by which Cys-loop receptor activity is regulated in relation to learning-dependent synaptic plasticity are not well-understood.

One way these questions can be approached is using anatomically-simple, genetically-tractable organisms such as the nematode *C. elegans. C. elegans* has a small nervous system consisting of 302 neurons, whose connectivity has been completely mapped (White et al., 1986). Remarkably, the *C. elegans* genome contains over 100 different Cys-loop LGC genes, more than double the number found in the human genome. These include multiple nicotinic-like acetylcholine-gated cation channels and GABA_A_-like GABA-gated anion channels, as well as anion channels gated by glutamate (e.g. AVR-14), acetylcholine (e.g. ACC-1) and serotonin (MOD-1) (Hobert, 2013). Nematode LGCs have also been identified with novel ligands not known to be fast neurotransmitters in other animals, such as tyramine (Pirri et al., 2009; Ringstad et al., 2009) and dopamine (Ringstad et al., 2009). For many of these nematode channels, the activating ligand is not known, and basic channel properties have not been characterised.

Despite its small nervous system, *C. elegans* is capable of performing both non-associative and associative learning, which offers an opportunity to test the function of these LGCs in neural plasticity. For example, animals infected with pathogenic strains of bacteria learn to avoid their odourants, which are attractive to naive animals (Zhang et al., 2005).

Neuronal ablation and genetic experiments indicate that learned avoidance requires a neural pathway involving serotonergic chemosensory neurons called ADF (Ha et al., 2010; Zhang et al., 2005). ADF mainly synapse onto the interneurons AIZ and RIA, both of which play critical roles in the neural circuit underlying olfactory response to bacterial odorants and learning (Ha et al., 2010; Liu et al., 2018). Previous work show that the function of a worm homolog of 5-HT_3_, *mod-1*, in AIZ is required for the aversive learning (Jin et al., 2016; Zhang et al., 2005). However, it is not clear whether the synaptic signalling between ADF and RIA is important for learning. In particular, it is unknown whether a serotonin signal is important for the role of RIA in learning and what receptor may mediate this function of neural plasticity.

Here we describe a new serotonin-gated LGC from *C. elegans*, LGC-50, that plays a key role in the aversive olfactory learning. LGC-50 is one of five new monoamine-activated Cys-loop LGCs that we have identified in a systematic deorphanisation of previously uncharacterised channels. Like other monoamine-gated LGCs, LGC-50 is expressed in neurons postsynaptic to aminergic neurons, specifically in the RIA neurons known to be critical for serotonin-dependent pathogen avoidance learning. Interestingly, LGC-50, is required for learned pathogen avoidance, and its regulated expression appears to be an important component in the plasticity mechanism.

## Results

### De-orphanisation of new amine-gated LGCs

Although a number of monoamine receptors, metabotropic as well as ionotropic, have been identified in *C. elegans*, many neurons receiving synaptic input from aminergic neurons express no known aminergic receptor. We reasoned therefore that some of the uncharacterised Cys-loop LGCs might be receptors for monoamine neurotransmitters. Three *C. elegans* LGCs have been previously described as monoamine-gated (Pirri et al., 2009; Ranganathan et al., 2000; Ringstad et al., 2009), but many predicted LGCs in the worm genome, including several closely-related channels, had no characterised endogenous ligand. A phylogenetic analysis of 171 *C. elegans* LGC genes (Table S1), based the analysis on the entire gene sequences, revealed the presence of a subfamily (Fig. 1) including the known monoamine-gated LGCs along with several uncharacterised channels. In line with previous reports we found that within this putative monoamine-gated subfamily (Fig.1 highlighted), the *mod-1* and *lgc-50* genes diverge from other genes in the group (Jones and Sattelle, 2008).

**Figure 1.**
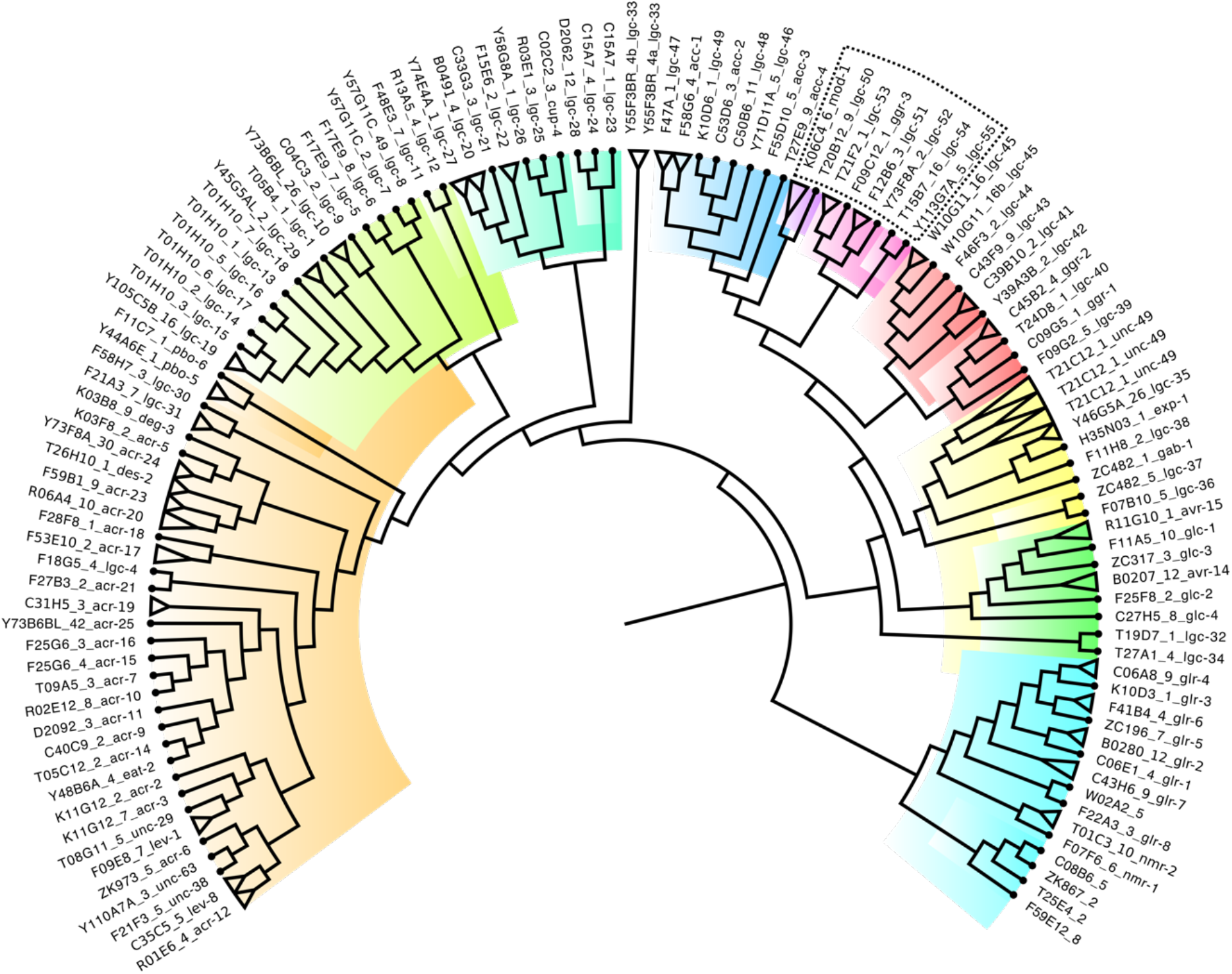
The superfamily of ligand-gated ion channel genes of C. elegans. Phylogenetic tree of the ligand-gated ion channel genes of C. elegans divided by colours into subfamilies and with the predicted amine-gated group highlighted in the dotted box. From left to right: orange, green and turquoise highlight subfamilies of the cationic nicotinic acetylcholine-like group of channels. Blue highlights the ACC group of acetylcholine-gated anion channels, purple the serotonin-gated channels, pink the amine-gated group, yellow the GABA-gated group, green the glutamate-gated anionic group and light blue the glutamate-gated cationic group. Where isoforms are present, these have been collapsed and are represented as triangles. Dotted lines highlight the aminergic subgroup.

We next initiated a systematic screen to deorphanise *C. elegans* LGCs without known ligands. Initially, we generated cDNA clones of the 5 orphan LGC genes in the putative monoamine-gated group: *lgc-50, lgc-51, lgc-52, lgc-54*, and *ggr-3*. By heterologous overexpression in *Xenopus* oocytes and two-electrode voltage clamp recordings we measured potential channel activity evoked by application of a panel of monoamine and other neurotransmitters. We surveyed responses to 11 putative agonists, including all monoamines known to be used in *C. elegans* (dopamine, serotonin, tyramine and octopamine), classical neurotransmitters used by *C. elegan*s (acetylcholine, GABA and glutamate) and other potential neurotransmitters and neuromodulators (betaine, tryptamine, histamine and glycine). All possible ligands were screened at 100 µM, a concentration well above the expected EC50 value.

In this way, we identified ligands for four of the five LGCs in this group. Three of the receptors, GGR-3, LGC-52 and LGC-54, were activated by both dopamine and tyramine (Fig. 2C-E); one of these, GGR-3 also displayed a small response to octopamine at very high (1 mM) concentrations (Fig. 2 & Supplementary Fig. S1A). Of these three receptors, LGC-52 showed a clear ligand preference for dopamine: its EC_50_ value was a 100-fold lower for dopamine than for tyramine, and dopamine also evoked a larger peak current. In contrast, although both GGR-3 and LGC-54 exhibited higher sensitivity to tyramine over dopamine, as shown by a significantly lower EC_50_ (inserts fig. 2D-E), they showed larger responses to dopamine. In contrast, we observed LGC-50 to be gated by 5-HT, with an EC_50_ of 0.94 µM (Fig. 2F), as well as the 5-HT metabolite tryptamine (Fig. 2A). The final orphan gene of this subfamily, LGC-51, did not exhibit any specific currents in response to the ligands tested here when expressed alone, though the protein appeared to be expressed in the oocytes due to failing viability 4 days after injection as compared to water injected controls (Supplementary Fig. S1B).

**Figure 2.**
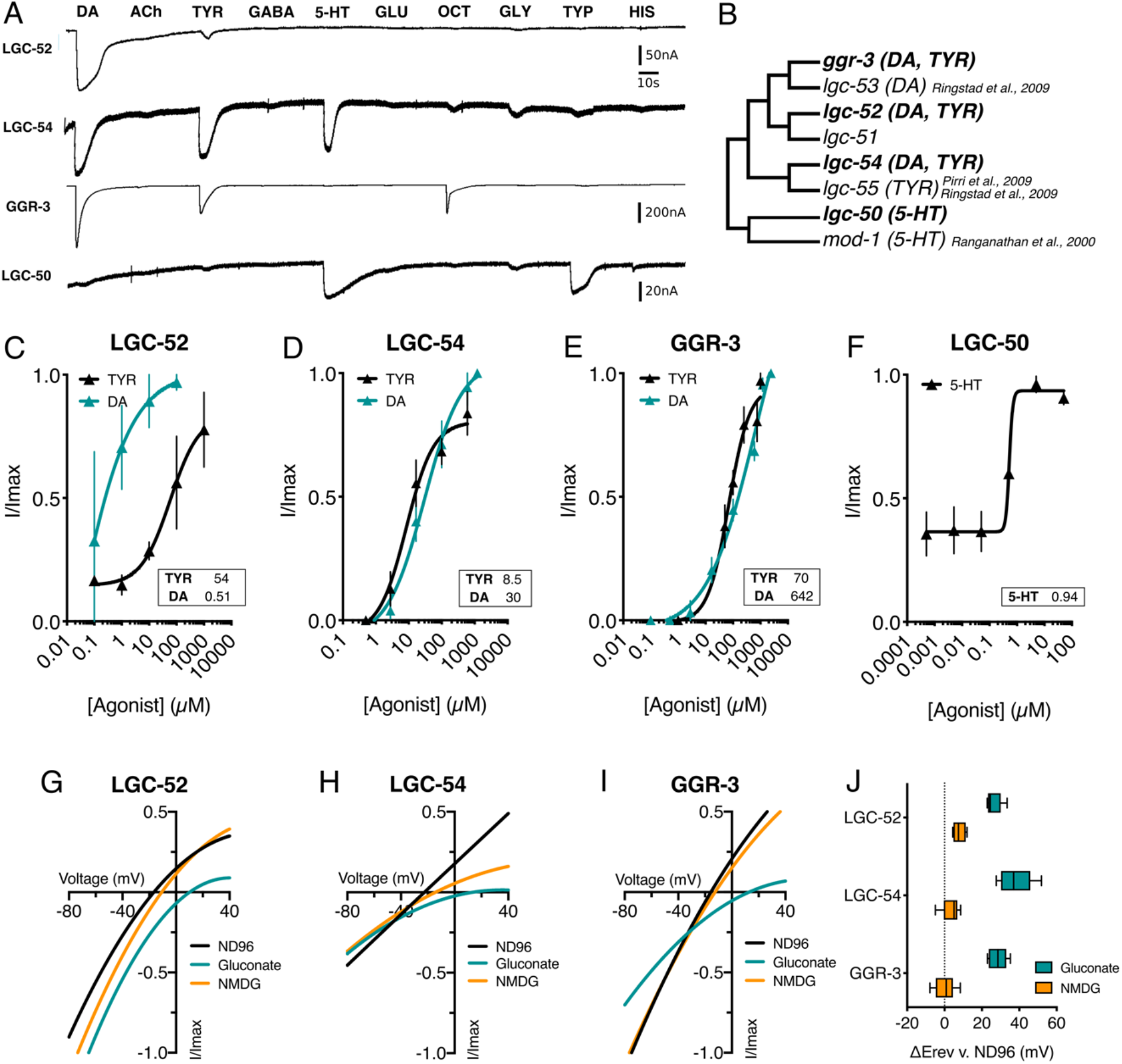
De-orphanisation of ligand-gated ion channels. A. Representative traces of continuous recordings of Xenopus oocytes clamped at -60 mV expressing LGC-52, LGC-54, GGR-3 and LGC-50, exposed to 100 µM DA, ACh, TYR, 5-HT, GLU, OCT, GLY, TYP & HIS. B. The predicted aminergic group based on phylogenetic analysis (Fig 1), showing orphan and characterised amine gated LGCs, highlighting lgc-50 and mod-1 as separated from the predicted dopamine-gated channels. C-F. Agonist-evoked dose response curves from oocytes expressing aminergic LGCs. Curves fitted to the Hill equation with variable slope using current normalised to I_max_ for each oocyte. Inserts show EC_50_ in µM. Error bars represent SEM for 4-9 oocytes. G-I. Representative current-voltage relationships for oocytes expressing LGC-52, LGC-54 and GGR-3. The current was normalised to I_max_ for each oocyte and baseline current subtracted from agonist-evoked current, with agonist present at EC_50_ concentrations. J. Average calculated from 4-10 oocytes for each construct of ΔE_rev_ in NMDG or Gluconate vs. in ND96 displays a positive shift of E_rev_ in gluconate indicating preference for anions. DA: dopamine, ACh: acetylcholine, TYR: tyramine, 5-HT: serotonin, GLU: glutamate, OCT: octopamine, GLY: glycine, TYP: tryptamine, HIS: histamine.

We next investigated the ion selectivity of the newly deorphanised channels. We carried out ion substitution experiments in oocytes expressing nematode LGCs, with the shift in reversal potential (Δ E_rev_) for a Na^+^ free (NMDG) or low Cl^-^ (Na Gluconate) solution compared to a solution with high sodium and chloride (Fig. 2G-I). For oocytes expressing each of the three channels, GGR-3, LGC-52 and LGC-54, the average shift in low Cl^-^ solution was significantly larger than the shift in Na^+^ free solution, with a mean shift in low Cl^-^ solution of 26-37mV (Fig. 2J). These results indicate that all three are inhibitory, anion-selective channels. Interestingly, the anion selectivity of previously-described pentameric LGCs depends in part on the conserved PAR motif in the M2-3 intracellular loop (Jensen et al., 2005), which is thought to line the channel pore and act as a pore gate (Laverty et al., 2019). All three of the newly deorphanised anion-selective receptors, LGC-52, LGC-54 and GGR-3, contain a PAR motif (Supplementary Fig. S2A).

### Aminergic LGCs are expressed postsynaptically to aminergic neurons and identification of a heteromeric LGC

Many of the principal synaptic targets of aminergic neurons have not been reported to express aminergic receptors (Bentley et al., 2016). We therefore speculated that these newly deorphanised aminergic channels might be expressed in some of these neurons and thereby mediate aminergic synaptic transmission. To address this question, we determined the expression pattern of each gene using fluorescent transcriptional reporters, containing both the upstream promoter region and the genomic DNA of each gene. We found that the 5-HT gated channel *lgc-50*, was strongly expressed in the RIA neurons (Fig. 3A). RIA is one of the two principal post-synaptic targets of ADFs, a pair of serotonergic neurons in the head. Interestingly, the other major synaptic target of the ADFs are the AIZs, which have been shown previously to express the other *C. elegans* serotonin-gated channel, *mod-1* (Gürel et al., 2012). We did not obverse any overlap in expression of *lgc-50* and *mod-1*, suggesting they have distinct and separate roles in serotonergic communication. These results are consistent with serotonin-gated channels playing a key role in fast serotonergic neurotransmission.

**Figure 3.**
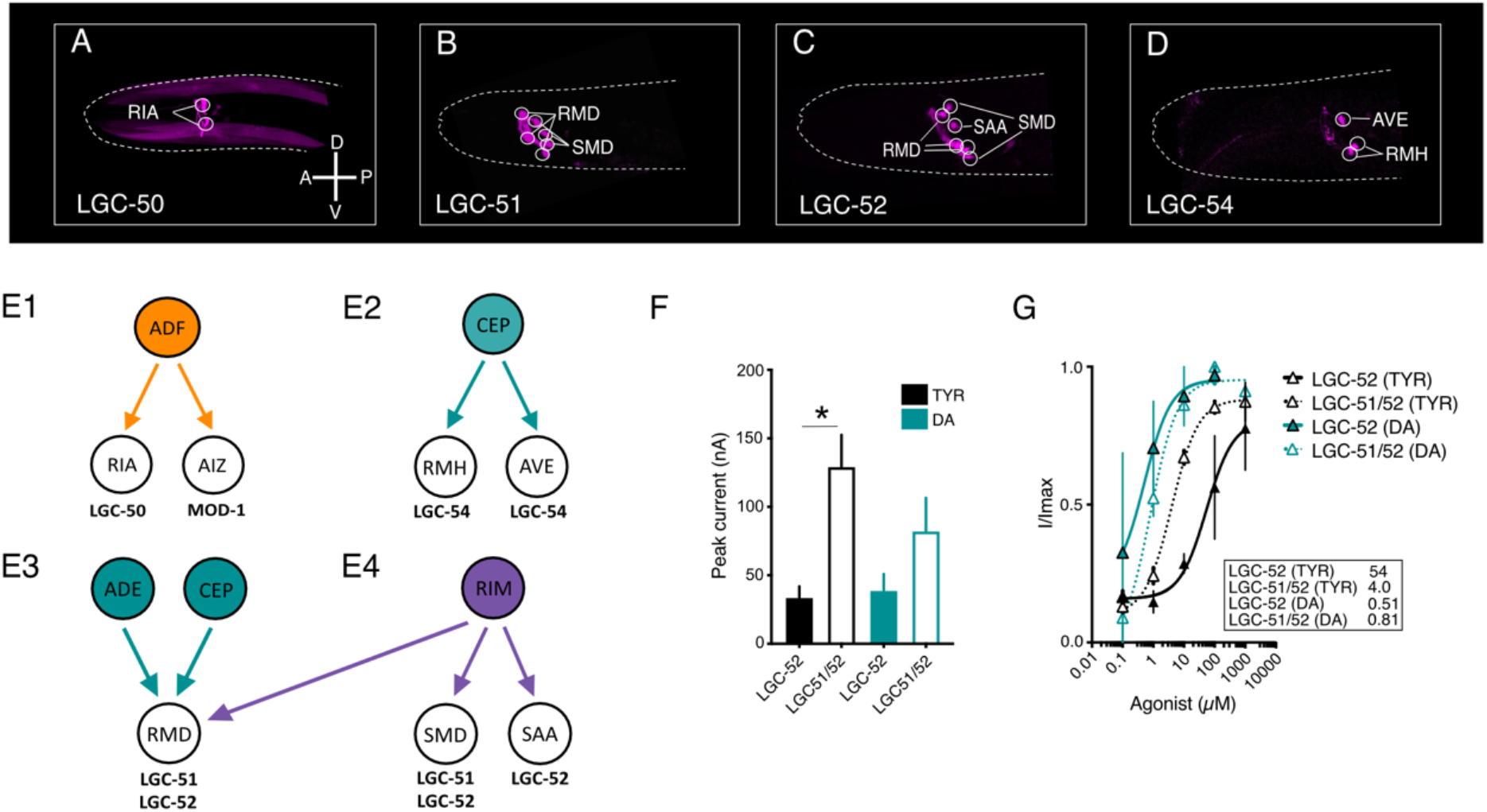
Novel amine gated LGCs are expressed in major synaptic targets of aminergic neurons and identification of a heteromeric LGC. A-E. Reporter lines expressing intercistonically spliced mKate or GFP under gene specific promotors reveal that all newly identified aminergic LGCs are localised in neurons that to some extent are postsynaptic to aminergic neurons (e.g. ADF, ADE, CEP, RIM). A. Plgc-50:lgc-50 gDNA:SL2 mKate expression was observed in RIA as well as head muscles. B. Plgc-51:lgc-51 gDNA:SL2 GFP expression was observed in RMD and SMD. C. Plgc-52:lgc-52 gDNA:SL2 GFP expression was observed in SMD, RMD and SAA. D. Plgc-54:lgc-54 gDNA:SL2 mKate was found to be expressed in AVE and RMH. E1-4. Graphical representation of neurons postsynaptic to aminergic neurons that express newly identified aminergic LGCs. F. Oocyte peak current (nA) in response to application of 1mM TYR or DA showing significantly higher peak currents in the LGC-51/52 heteromer compared to LGC-51 homomer. Bars represent SEM of 5-14 repeats. G. Agonist-evoked dose response curves from oocytes expressing monomeric LGC-51 or heteromeric LGC-51/52. Curves fitted to the Hill equation with variable slope using current normalised to I_max_ for each oocyte. Inserts show EC_50_ in µM for DA and TYR. Error bars represent SEM for 4-9 oocytes Scale bar indicates 100 µm. * P<0.05 vs. LGC-52 (TYR) by ordinary one-way ANOVA with Tukey’s multiple comparison correction.

For dopamine and tyramine gated channels we likewise observed expression in many neurons postsynaptic to dopaminergic and tyraminergic neurons. For example, *lgc-54* was strongly expressed in two neuron pairs AVE and RMH (Fig. 3D). Both these neurons are major postsynaptic targets of the dopaminergic CEP neurons (White et al., 1986), and neither neuron has been reported to express previously-described dopamine receptors. Likewise, clear *lgc-52* expression was observed in the RMD, SMD, and SAA neurons (Fig. 3C); all these neurons are major synaptic targets of the tyraminergic RIMs, and the RMDs are additionally targets of the dopaminergic CEP and ADE neurons (Fig. 3F 1-2). The expression pattern for *ggr-3* was much broader as compared to the other dopamine-gated channels, and we identified expression in BAG and ASH neurons as well as a number of yet unidentified neurons (Fig. S3).

Interestingly, the reporter for *lgc-51*, the only channel with no ligand response in our *Xenopus* oocyte screen, also expressed specifically in the RMD and SMD neurons (Fig. 3B), both of which express its most closely related paralogue *lgc-52*. We therefore hypothesised that LGC-51 and LGC-52 might form functional heteromers in these cells. To address this question, we co-expressed LGC-51 and LGC-52 in *Xenopus* oocytes and compared the responses to oocytes expressing LGC-52 alone. We indeed observed currents in the LGC-51/52 expressing oocytes that were distinct (Fig. 3F-G) from currents observed in oocytes expressing LGC-52 alone. For example, while the EC_50_ for dopamine was similar between the LGC-52 homomer and the LGC-51/52 heteromer (at 0.51 µM and 0.81 µM respectively), LGC-51/52 heteromers showed a much higher potency for tyramine (4 µM EC_50_) compared to the LGC-52 homomer (54 µM Fig. 3G). Likewise, the tyramine-induced peak current achieved by oocytes expressing the LGC-51/52 heteromer was also significantly higher than those expressing LGC-52 alone, although there was no significant effect on the dopamine peak current. Together, these results suggest that heteromerisation of LGC-51 with LGC-52 predominately effects its gating efficiency by tyramine (Fig. 3F), changing a channel with a strong preference for dopamine to one that is effectively activated by both tyramine and dopamine.

### Dopamine and tyramine gated channels vary in their response to repeated stimulation and antagonist application

We sought to better understand the individual properties of these seemingly similar dopamine and tyramine gated LGC channels. To do so we recorded the recovery time of the channels by exposing oocytes to multiple pulses of each agonist with varying pulse intervals, during which time the oocyte was perfused with ND96 buffer (Fig. 4). For LGC-50 and MOD-1, both gated by 5-HT there was no significant difference in recovery time between the channels (Fig. 4A). In contrast, in response to multiple applications of dopamine, GGR-3 showed a significantly reduced peak current ratio at 10s and 30s intervals compared to LGC-54, LGC-52 and LGC-51/52, and did not recover to the maximal peak size until 60s after the initial pulse (Fig. 4B, E). At the 10s pulse interval LGC-52 also showed significantly slower recovery to initial pulse than LGC-54 and LGC-51/52 (Fig. 4B). Both LGC-54 and the LGC-52/51 heteromer, in contrast, showed no significant reduction in peak ratios following repeated stimulation. A similar trend was observed in response to multiple applications of tyramine, with only GGR-3 showing a significantly decreased peak ratio at 10s pulse interval as compared to the initial pulse (Fig. 4E-F).

**Figure 4.**
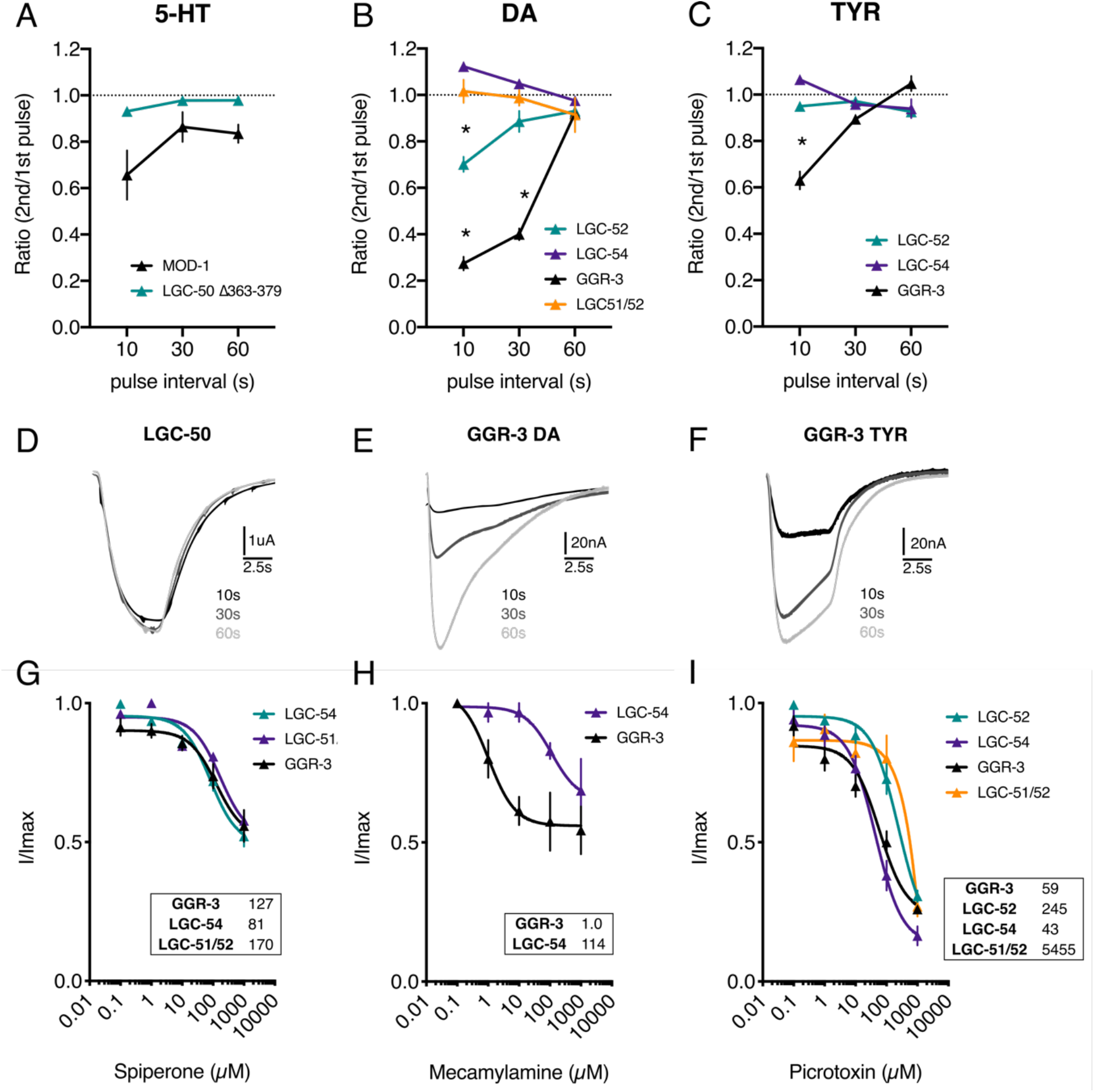
Differences in agonist occupancy and antagonistic profile for dopamine-gated channels. A. Oocyte peak current ratio of two 10s agonist pulses with varying pulse intervals (s) during which the oocyte is washed with ND96 buffer. Agonist was delivered at the EC_50_ concentration for each channel. Dashed line at ratio = 1. Error bars represent SEM of 3-8 repeats. *P<0.05 compared to all other constructs at the same pulse interval, calculated by 2way ANOVA with Tukey’s multiple comparison correction. B. Antagonistic inhibitory dose response curves from oocytes expressing aminergic LGCs. Agonist concentration remained constant throughout, at the respective EC_50_ for each channel. Curves fitted to the Hill equation with three parameter slope using current normalised to Imax for each oocyte. Inserts show IC_50_ in µM. Error bars represent SEM for 4-9 oocytes.

We further examined possible differences between these channels by investigation of the antagonist profiles of three potential antagonists; mecamylamine, spiperone and picrotoxin. These chosen antagonists are known to target different binding sites and have different specificity for vertebrate receptors. We found that mecamylamine, a nicotinic acetylcholine receptor blocker which is thought to act by binding in the ligand binding region, was able to partially block GGR-3 with an IC_50_ of 1 µM; in contrast only a partial block was achieved for LGC-54 with an IC_50_ of over 100 µM (Fig. 4H). This vast difference in IC_50_ suggests that despite binding the same aminergic ligands, the ligand binding domains of GGR-3 and LGC-54 may be structurally different. Spiperone, which preferentially binds dopaminergic binding sites in mammals (Zhen et al., 2010), led to a partial block of GGR-3, LGC-51/52 and LGC-54 with comparable IC_50_ values of 127 µM, 170 µM and 81µM respectively (Fig. 4G). In contrast, picrotoxin, a well-characterised anion pore blocker (Newland and Cull-Candy, 1992), led to a more complete inhibition of currents, in particular for GGR-3 and LGC-54 with IC_50_ values of 59 µM and 43 µM respectively. Interestingly, the IC_50_ of picrotoxin of the LGC-51/52 heteromer was an order of magnitude larger than that of the LGC-52 homomer (Fig. 4I). This suggests that the pore structure and size of the LGC-52/51 heteromeric channel differs significantly from the LGC-52 homomer.

### LGC-50 is a cationic channel whose trafficking is regulated by its large intracellular domain

In contrast to the new dopamine and tyramine receptors, the newly-deorphanised serotonin receptor LGC-50 was difficult to characterise in detail due to the small size of currents. The application of 5-HT to oocytes expressing LGC-50 elicited only small peak currents, on average 15nA (Fig. 2A), despite detection of high protein concentrations (Supplementary Fig S4); these currents were nonetheless dose dependent (Fig. 2F) and not present in control oocytes (Supplementary Fig. S5A). In contrast, oocytes expressing the other *C. elegans* ionotropic serotonin receptor MOD-1 (Ranganathan et al., 2000) displayed much larger currents only after 2 days of incubation, suggesting a higher degree of protein expression (Supplementary Fig. S5B). The large intracellular loop between transmembrane helices 3 and 4 is widely accepted to be involved in the trafficking and proper cellular localisation of ligand-gated ion channels (Lo et al., 2008; Perán et al., 2006). This domain is the most variable section of ligand-gated ion channels, and contains many protein-protein binding sites and sites for post translational modifications (Chen and Olsen, 2007). The two closely related serotonin-gated channels in *C. elegans, mod-1* and *lgc-50*, have a high degree of sequence identity of 47% outside of the M3/4 loop, however this falls to just 15% when only comparing the M3/4 loop of the two proteins (Supplementary Fig. S2B). Thus, we wondered whether the small currents observed in *lgc-50*-expressing oocytes might be a result of poor membrane localisation of LGC-50 protein due to regulatory domains within the M3/4 loop.

To investigate this possibility, we exchanged the intracellular M3/4 loop of LGC-50 with the equivalent region of MOD-1 (LGC-50:MOD-1 327-458) and expressed the chimeric receptor protein in oocytes. This resulted in a significant 175-fold increase in peak current relative to the native LGC-50; the peak current amplitude (2.6 µA) did not differ significantly to the peak current of wild type MOD-1 (2.9 µA; Fig. 5B). In the converse experiment, we exchanged the MOD-1 M3/4 loop for that of LGC-50 (MOD-1:LGC-50 325-462); this resulted in a significant 44-fold decrease in peak current to 66 nA, which in turn did not differ significantly to the peak current observed in wild type LGC-50 (Fig. 2B and Supplementary Fig. S5B). In each case the dose dependency and EC_50_ of the chimeric channels matched that of the recipient channel, not that of the donor M3/4 domain (Fig. 5C-D). This suggests that the change in peak current conferred by the M3/4 region was due to increased or decreased membrane surface localisation rather than changes to ligand binding efficacy or gating properties. Taken together this data suggests that the M3/4 loop of LGC-50 might contain domains that are able to restrict plasma membrane trafficking of the channel.

**Figure 5.**
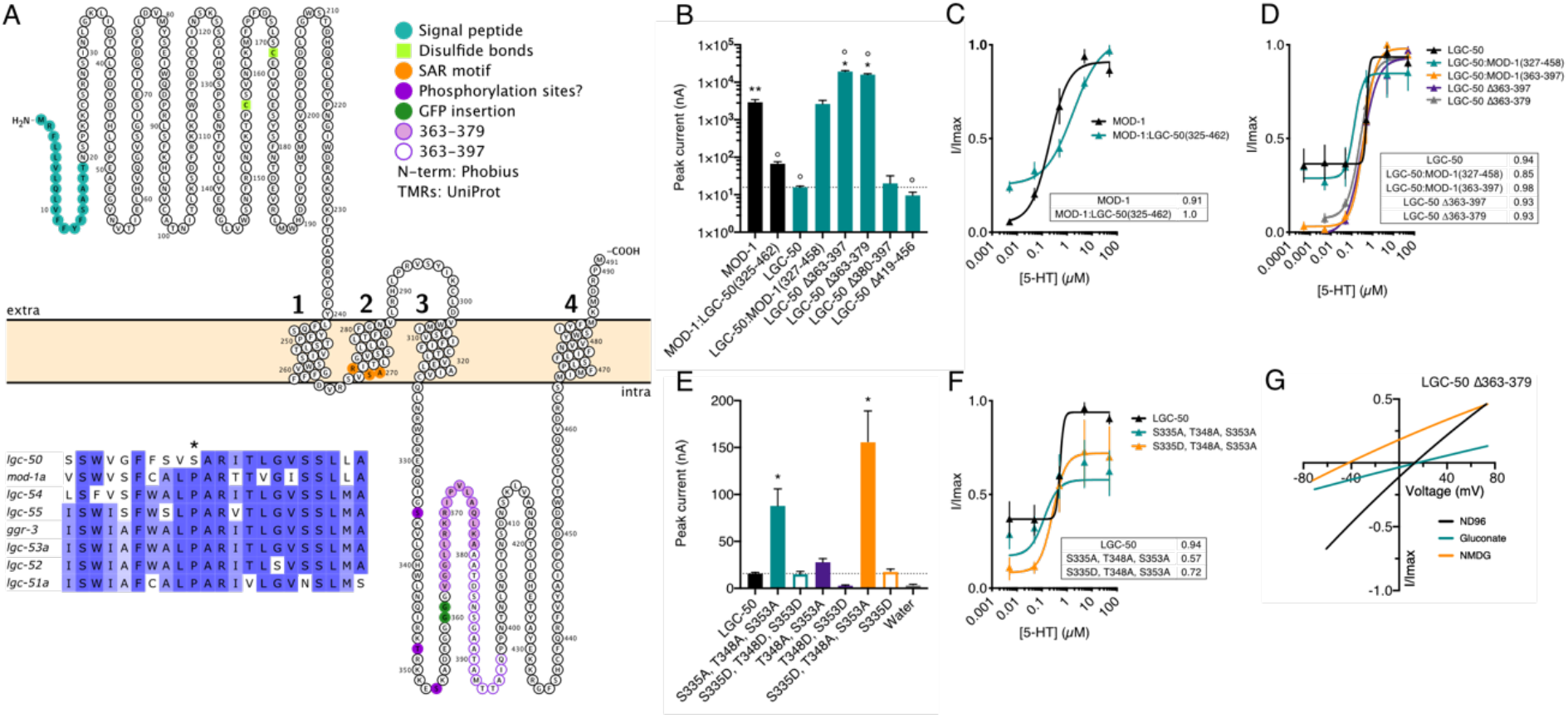
Identification of LGC-50 as a cationic channel and a binding motif for regulating surface localisation. A. Topology diagram of LGC-50 depicting the location of transmembrane and functional protein domains. Insert: alignment of the “PAR “motif of aminergic C. elegans LGCs showing the substitution of proline to serine in LGC-50. B. Oocyte peak current (nA) in response to application of 5 µM 5-HT on different chimera version of MOD-1 and LGC-50. Bar represents mean + SEM of 2-26 repeats. Dashed line positioned at the wild type LGC-50 peak current. * P<0.05 vs. LGC-50, ° P<0.05 vs. MOD-1 by ordinary one-way ANOVA with Dunnett multiple comparison correction C. 5-HT evoked dose response curves from oocytes expressing wild type MOD-1 and MOD-1 mutants. Error bars represent SEM of 8-12 oocytes. Curves fitted to the Hill equation with variable slope using current normalised to Imax for each oocyte. Insert shows EC_50_ in µM. D. 5-HT evoked dose response curves from oocytes expressing LGC-50 and MOD-1 mutants. Error bars represent SEM of 3-6 oocytes. Curves fitted to the Hill equation with variable slope using current normalised to Imax for each oocyte. Insert shows EC_50_ in µM. E. Oocyte peak current (nA) in response to application of 5 µM 5-HT on phosphomimic versions of LGC-50. Bar represents mean + SEM of 2-30 repeats. Dashed line positioned at the wild type LGC-50 peak current. * P<0.05 vs. LGC-50, by ordinary one-way ANOVA with Dunnett multiple comparison correction. F. 5-HT evoked dose response curves from oocytes expressing LGC-50 phosphomimic mutants. Error bars represent SEM of 3-6 oocytes. Curves fitted to the Hill equation with variable slope using current normalised to Imax for each oocyte. Insert shows EC_50_ in µM. G. Representative current-voltage relationships for oocytes expressing LGC-50 del363-379. Current was normalised to I_max_ for each oocyte and baseline current subtracted from agonist evoked current, with agonist present at EC_50_ concentration.

In order to identify such domains, we tested the effects of deletion mutations in the LGC-50 M3/4 loop. Several deletions of the LGC-50 M3/4 loop were generated, including Δ363-397, Δ363-379, Δ380-397 and Δ419-456 (depicted in Fig. 5A). Two of these deletions, in the latter half of the loop from 380-456, had no significant effect on the peak current when compared to wild type LGC-50 (Fig. 5B). However, two deletions–Δ363-397, and the smaller deletion within this region Δ363-379–resulted in significant increases in peak current to 19 µA and 15 µA respectively (Fig. 5B). Indeed, the currents of these LGC-50 deletion mutants significantly surpassed the peak current achieved by either the LGC-50::MOD-1 chimera or wild type MOD-1. Again, each of these deletion mutations showed similar dose dependency and efficacy of serotonin response to wild-type LGC-50 (Fig. 5D) as well as similar protein expression as wild-type LGC-50 (Supplementary Fig. S4), supporting the notion that these mutations alter cell surface trafficking rather than other properties of the channel. These data strongly suggest the presence of a functional domain within this 16 amino acid region that results in the severe restriction of cell surface trafficking.

In addition to the aforementioned 16 amino acid region, we also investigated three predicted phosphorylation sites upstream of this region. Phosphorylation of the M3/4 loop has previously been implicated in the trafficking and cell surface recycling of GABA_A_ receptors (Connolly et al., 1999; Jovanovic et al., 2004). The first site in the M3/4 loop of LGC-50, S335, is predicted to be a cdc2 site, and T348 and S353 are predicted PKC/PKA sites (Blom et al., 1999, 2004) (Table S2). We therefore introduced both phosphorylation-dead alanine mutations and phosphomimic aspartate mutations in order to understand whether phosphorylation at these residues may be important in receptor trafficking. We observed significant 5-fold and 10-fold increases in peak current amplitude for two mutation combinations; S335A, T348A, S353A and S335D, T348A, S353A respectively. Dose dependency was not affected by either of these mutations, again suggesting an effect of the number of receptors at the surface rather than other changes to channel properties (Fig. 5E-F).

As we were now able to induce efficient cell surface trafficking of LGC-50, we sought to determine the ion selectivity of the channel using the M3/M4 deletion mutants. We carried out ion substitution experiments as done previously with the dopamine/tyramine receptors, comparing the reversal potential shifts for sodium and chloride. Interestingly, we observed that LGC-50 Δ363-379 is selective for cations, with an average reversal potential shift in Na^+^-free solution (NMDG) of –43mV +/- 4mV (Fig. 5G). Thus, despite its phylogenetic proximity to GABA_A_ receptors and other anion-selective ligand-gated channels, *lgc-50* appears to encode an excitatory, cation selective serotonin receptor. It is worth noting that unlike all other members of the aminergic LGC group, LGC-50 does not contain the PAR region which typically confers anion selectivity (Jensen et al., 2005) (Fig. 5A alignment insert); instead, the proline residue thought to be important for ion selectivity through the pore is substituted by a serine residue.

To investigate the localisation of LGC-50 *in vivo*, we used transgenic animals expressing a GFP-tagged LGC-50 protein from the *lgc-50* locus. GFP-tagging did not appear to compromise the receptor’s ability to functionally traffic to the plasma membrane, as expression of a GFP tagged version of the receptor carrying the Δ363-379 deletion in oocytes generated currents of comparable size to the untagged receptor (Fig. 6E). When we cultured the transgenic animals on E. coli, we observed little expression of tagged LGC-50 in the nerve ring, the site of most neuronal processes (Figure 6A-B). However, when we exposed worms to the pathogenic bacteria *Serratia marcescens*, we observed visible nerve ring expression of LGC-50::GFP in a punctate pattern (Fig. 6C-D) This result indicates that the control of LGC-50 expression that we observe in oocytes may be used in vivo to regulate receptor abundance at synapses.

**Figure 6.**
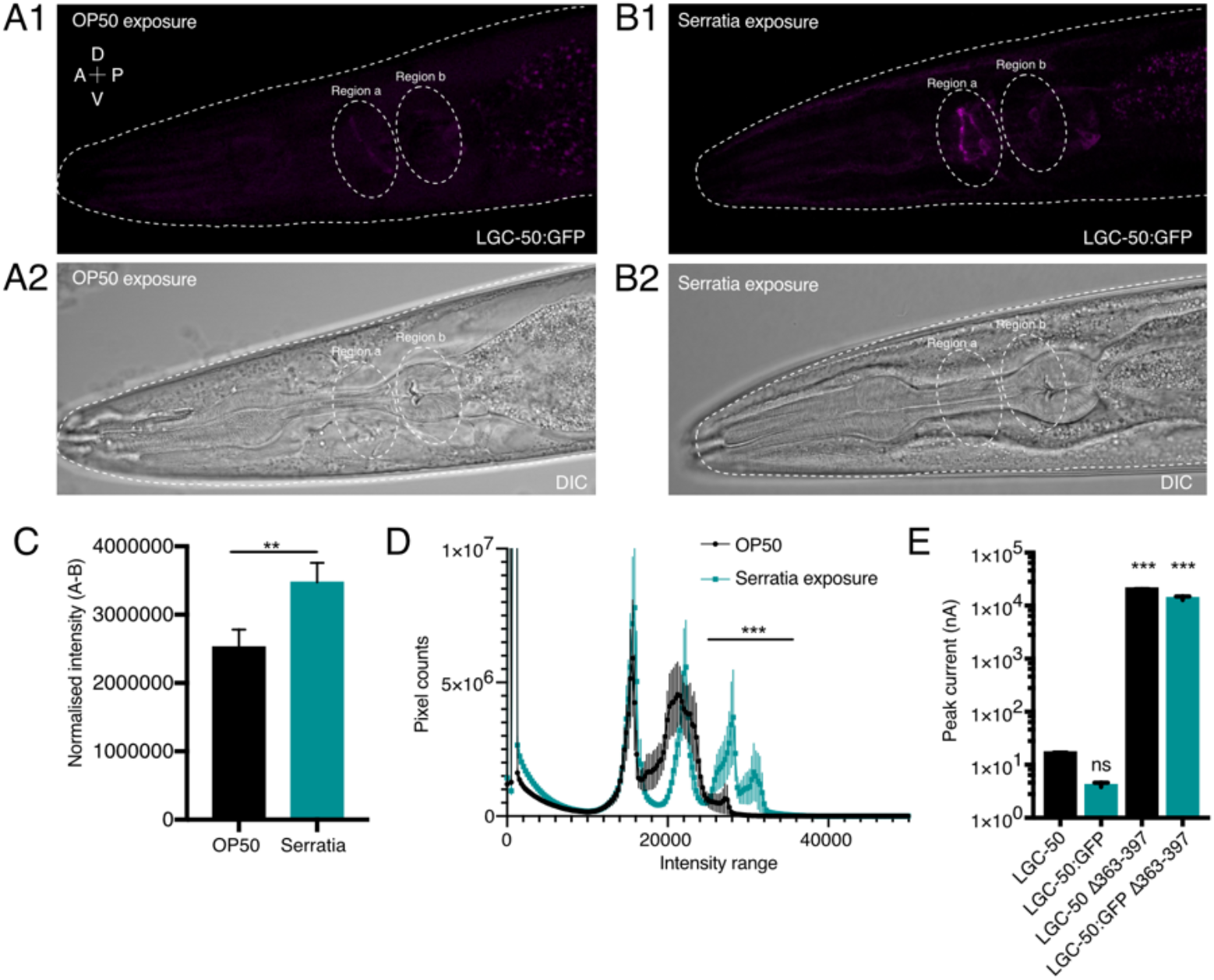
A1-2. Representative image of an OP50 exposed worm annotating the two regions A and B that was used for intensity measurements. B1-2. Representative image of a Serratia Marcescens exposed worm annotating the two regions A and B that was used for intensity measurements. C. Normalised fluorescent intensity for region A subtracting background levels from region B indicate that the nerve ring intensity in Serratia exposed animals is significantly higher than after OP50 exposure. Bar represents mean + SEM. D. Whole worm intensity histogram displaying a right shift in pixel counts towards a significantly higher intensity range for Serratia exposed worms, indicating a more punctate distribution of LGC-50 protein., n= 20 OP50, 21 Serratia. E. Oocyte peak current (nA) in response to application of 50 µM 5-HT in lgc-50 wt or GFP tagged lgc-50 showing that GFP tagging do not influence the function of the receptor and that the Δ363-379 deletion still allows the receptor to traffic to the membrane. Bar represents mean + SEM of 4-26 repeats. **P<0.01 *** P<0.001

### LGC-50 in RIA is involved in aversive olfactory learning

LGC-50 shows specific and prominent expression in the RIAs, a pair of interneurons receiving synaptic input from the serotonergic ADFs. Naive *C. elegans* show strong attraction to odours produced by pathogenic *Pseudomonas aeruginosa* (PA14) bacteria; however, animals that have been exposed to and infected by pathogen learn to reduce their preference for these odours. The ADF-RIA synapses represent the first step in the learned aversion pathway defined by cell ablation experiments (Ha et al., 2010), and serotonin-deficient mutants likewise are defective in learned pathogen avoidance (Jin et al., 2016; Liu et al., 2018; Zhang et al., 2005). We therefore wondered whether LGC-50 might play an important role in aversive olfactory learning of pathogenic bacteria.

To investigate this question, we investigated the effect of an *lgc-50* deletion mutation on aversive learning. We used a previously established chemotaxis assay to measure the steering movement towards the odorants of PA14 (Ha et al., 2010; Liu et al., 2018). The efficiency of olfactory chemotaxis was calculated using the navigation index (Fig. 7A), as well as the total travelling distance (Fig. 7B-C, Materials and Methods). Consistent with our previous findings, after training with PA14 wild-type worms reduced the navigation index in the chemotactic steering towards PA14 odorants and travelled a significantly longer distance before reaching PA14. In contrast, *lgc-50* null mutants showed no difference in navigation index or in travelling distance after training with PA14 (Fig. 7B-C), indicating an olfactory learning deficiency for *lgc-50* mutants. Expressing a wild type *lgc-50* gene selectively in RIA fully rescued the learning defects measured by both navigation index and traveling distance, indicating that LGC-50 functions in the RIA neurons to promote aversive learning (Fig. 7D-E). In addition, the *lgc-50* mutants display wild-type chemotaxis towards the odorants of *E. coli* OP50, the standard food source for *C. elegans*, under both naive and training conditions (Fig. 7G-H), indicating intact chemotaxis ability. We also first used a two-choice assay (Liu et al., 2018; Zhang et al., 2005) which measured the olfactory preference of naive or trained worms for *P. aeruginosa* strain PA14 compared to *E. coli* OP50 and similarly identified a robust learning defect in *lgc-50* deletion mutant animals in comparison with wild-type controls (Fig. 7F). Furthermore, we tested *lgc-50* mutant worms on a separate learning paradigm – thermotaxis (Luo et al., 2014; Mori and Ohshima, 1995) and found no difference between wild type and *lgc-50* mutants (Supplementary Fig. S6), which demonstrate normal sensorimotor activity of *lgc-50* mutant animals. Together, these results demonstrate that LGC-50 in RIA regulates aversive olfactory learning.

**Figure 7.**
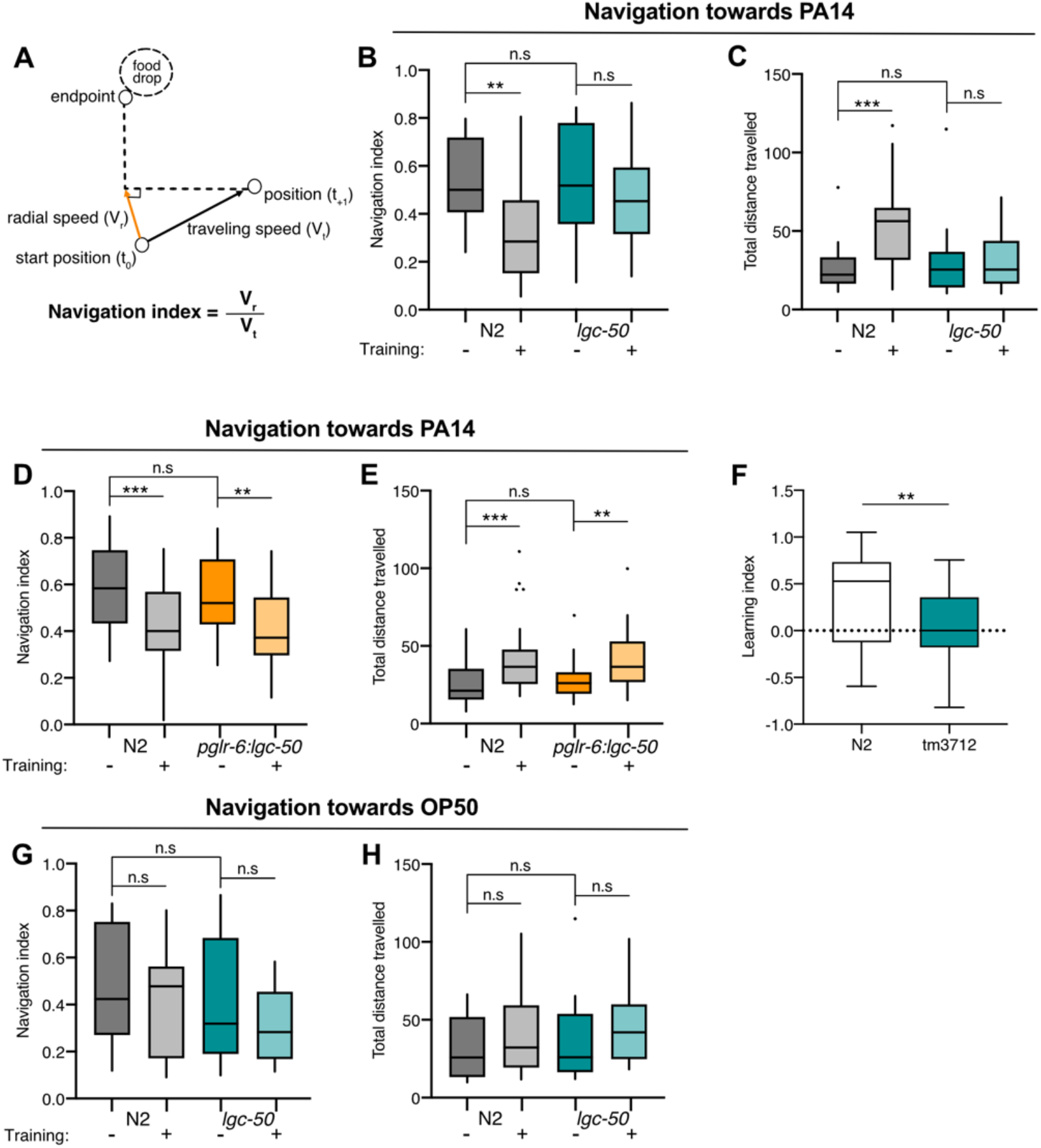
LGC-50 in RIA has a role in aversive olfactory learning. N2 animals and lgc-50 mutants were tested in the olfactory steering assay (B-E, G, H) or two-choice preference assay (F) after exposure to either OP50 or PA14. A. Schematics demonstrating the principle behind calculation of a navigation index from the olfactory steering assay. B-C. N2 animals trained on PA14 displayed a significantly decreased navigation index, together with an increased distance travelled to the food drop. Whereas lgc-50 mutants did not display any training-induced change in either navigation index or travelling distance. D-E. By reintroducing lgc-50 in RIA under the RIA-specific promotor glr-6 the learning deficit in the lgc-50 mutant was completely rescued. F. N2 and lgc-50 mutants significantly differed in the two-choice learning assay. N2: n=31, tm3712: n=41. Student’s t-test, ** P<0.01. G-H. The lgc-50 mutants are comparable to wild-type animals in their olfactory steering towards OP50 before and after training. For B-E, G-H: N2: n= 42 naïve animal, n= 38 trained animals, lgc-50 mutant: n= 40 naïve animal, n= 39 trained animals, pglr-6::lgc-50: n= 35 naïve animal, n= 42 trained animals. One-way ANOVA, ** P<0.01 *** P<0.001, boxes show first and third quartile, median and the whiskers extend to data points that are equal or less then 1.5 IQR from the quartiles. Dots represent outliers.

## Discussion

### Divergent channel properties of C. elegans monoaminergic LGCs

In this study, we describe five new LGCs activated by monoamines. We identified four channels that are gated by tyramine and dopamine, three of which are able to form functional homomers, GGR-3, LGC-54, LGC-52, as well as one heteromer consisting of LGC-51 and LGC-52. All four of these channels are anion selective; in contrast, the newly characterised serotonin-gated channel, LGC-50 is cation selective. These results reveal a remarkable evolutionary plasticity in the fundamental properties of ligand-gated channels. For example, LGC-54 shows its largest peak responses in physiological ranges to dopamine, yet its phylogenetically-closest relative LGC-55 forms a tyramine-selective channel. Likewise, the closest paralogue of GGR-3, a channel with highest potency achieved by tyramine, is LGC-53, a dopamine-selective channel. Closely related channels can also show divergent ion selectivity; the closest paralogue of the serotonin-gated cation channel LGC-50 is MOD-1, a serotonin-gated anion channel. Thus, the *C. elegans* monoamine-gated LGC subfamily shows remarkable diversification in the most fundamental properties of ligand binding and ion selectivity. *C. elegans* as well as many other invertebrate species contain a number of uncharacterised LGC families; it is interesting to speculate that these also may have evolved novel functional properties, possibly including novel activating ligands.

Even channels with nominally-similar ligand-binding profiles show interesting divergence in their expression pattern, kinetics and ligand preference. For example, amongst the newly identified dopamine and tyramine-gated channels, dopamine activated the homomeric LGC-52 channel most potently as compared to the other family members; however at EC50 concentrations dopamine elicited significantly larger peak currents for GGR-3 than LGC-52. This suggests that receptors may be differently localised *in vivo*, and that low affinity receptors such as GGR-3, requiring higher dopamine concentrations for activation, may be localised closer to agonist release sites in the active zones of the synapses and may evoke larger synaptic currents than higher-affinity receptors at lower dopamine concentrations possibly localised extrasynaptic. This phenomenon has been shown for the low affinity glutamatergic AMPA receptors and nAChRs (Biederer et al., 2017; d’Incamps and Ascher, 2014; Luo et al., 2014; Raghavachari and Lisman, 2004; Tang et al., 2016). In addition, we also observed differences in the capability of these receptors to undergo repeated stimulation by the same agonist. Repeated activation of GGR-3 led to a significant decrement in response magnitude, with the receptor only recovering full activity after a minute rest; in contrast there was no detectable wear-down or desensitisation of the other tyramine/dopamine receptors such as LGC-52. We also observed differences in the antagonist binding profiles of these channels, which may suggest structural differences in the ligand binding domains of these channels or differences in pore size between homomeric and heteromeric channels. In the future, the natural functional diversity of this ion channel family should provide a useful test bed to explore the relationship between LGC structure and function.

### Regulated trafficking and localisation of LGC-50 channels

Strikingly, our analysis of serotonin-gated LGCs has provided mechanistic insight into the regulation of LGC membrane trafficking and synaptic localisation. The serotonin-gated cation channel, LGC-50, when heterologously expressed in *Xenopus* oocytes, shows limited membrane expression in its native form, whereas the closely-related serotonin-gated anion channel MOD-1 shows robust constitutive trafficking (Ranganathan et al., 2000). Reciprocal domain swap experiments demonstrated that this difference in trafficking efficiency is specified by the cytoplasmic loop between the third and fourth transmembrane domains. We identified a domain of 17 amino acids in the intracellular M3/4 loop critical for controlling plasma membrane expression; when these residues were removed, the membrane expression of the receptor increased significantly, while dose dependency and ion selectivity were unaffected. Interestingly, both human GABAAR ß subunits and glycine receptors, which show significant homology to LGC-50 (25.84% with ß1; 27% for GlyR) appear to use similar molecular mechanisms to regulate cell surface localisation and trafficking. In both cases, the M3/4 loop has been strongly implicated in regulating trafficking, assembly and localisation of the receptors (Mele et al., 2016; Specht et al., 2011).

We also observed a potential role for phosphorylation in the fine control of LGC-50 plasma membrane expression. Specifically, we found that preventing phosphorylation of two predicted PKC sites in the M3/4 loop of LGC-50 led to significant increases in serotonin induced current without affecting dose dependency, whereas phosphomimic mutations led to a reduction. Again this parallels previous work on GABA_A_ receptors (Abramian et al., 2010; Chapell et al., 1998; Connolly et al., 1999; Nakamura et al., 2015) showing that phosphorylation of sites within the M3/4 loop of GABAAR by PKC induces receptor internalisation and plays major roles in synaptic plasticity at GABAergic synapses (Mele et al., 2016). Using peak current amplitude as a measure of the number of channels at the surface, the alterations made to these phosphorylation sites had a modest effect on trafficking compared to the previously describe deletion; however, multiple mechanisms may regulate cell surface expression and localisation *in vivo*. We hypothesise that the 16 amino acid region that we identified restricts surface expression, whereas regulation of the phosphorylation pattern may affect internalisation of the channels as it does in related channels (Connolly et al., 1999; Jovanovic et al., 2004).

Taken together, we observe a strikingly equivalent set of molecular mechanisms controlling the expression and membrane localisation of an excitatory serotonin-gated channel in *C. elegans* to those controlling the membrane localisation of inhibitory glycine and GABA_A_ receptors in mammals. This suggests a potentially high degree of mechanistic conservation in the regulation of LGC trafficking across diverse phyla and receptor type, which could be adapted in different circuits to generate neural and behavioural plasticity.

### An excitatory serotonin-gated LGC plays a critical role in associative learning

We have also identified a key role for LGC-50 in aversive learning and memory. Previous work demonstrated that learned avoidance of odours given off by pathogenic bacteria following infection depends on serotonin and on the RIA interneurons, which receive extensive serotonergic innervation (Ha et al., 2010). We find that *lgc-50* mutants are defective in pathogen avoidance learning, though their initial responses to pathogen odours are normal. This learning defect could be rescued by cell-specific expression of *lgc-50* in the RIA neurons, indicating that LGC-50 channels function in the RIA neurons to facilitate learned aversion. Interestingly, we observed that exposure to a different pathogen regulates the expression of LGC-50 channels; whereas little expression of LGC-50 was found in the nerve ring under normal growth conditions, its abundance was strongly enhanced following infection with pathogenic bacteria. Thus, we speculate that learning-induced expression of LGC-50 could enhance the localization of the serotonin-gated channel at the synapse, which underlies the critical role of LGC-50 in learning.

How might LGC-50’s activity remodel the olfactory circuit to alter odorant preferences following pathogen exposure? Previous work has indicated that learned pathogen aversion depends specifically on the serotonergic ADF neurons and their synaptic targets the RIA interneurons. Functional analyses on RIA suggest that following training, this pathway acts to inhibit steering and promote turning in the presence of pathogen odours through RIA synapses onto the SMD motorneurons (Ha et al., 2010; Liu et al., 2018). Our results suggest that induced expression of LGC-50 in the process of RIA, together with training-regulated serotonin signal (Qin et al., 2013; Zhang et al., 2005) is an important mechanism for the mobilisation of this aversive pathway following training. Previous work has shown that another worm homolog of 5-HT3 receptors, MOD-1, acts in a different group of interneurons to regulate aversive olfactory learning (Zhang et al., 2005). With the results on LGC-50, these findings together highlight the critical role of serotonin-gated channels in neural plasticity.

The results described here may provide more general insight into the roles of pentameric LGCs in learning and memory. In particular, mammalian 5-HT3 receptors, which like LGC-50 are serotonin-gated cation channels, have been implicated in various forms of learning and behavioural plasticity. For example, 5-HT_3_ receptors have been shown to play a key role in reward pathways, with their insertion at synapses between the dorsal raphe nuclei and the VTA, by thus promoting enhanced dopamine release (Wang et al., 2019). Moreover, regulation of 5-HT_3_ receptor expression and abundance has been shown to be important for fear extinction (Kondo et al., 2014). In addition, changes in the expression of 5-HT_3A_ receptors in the mouse visual cortex are important for cross-modal plasticity following sensory loss (Lombaert et al., 2018). The molecular mechanisms by which 5-HT_3_ receptor activity is regulated to generate synaptic plasticity in these examples are currently not well-understood. In the future, it will be interesting to investigate whether regulated trafficking mechanisms similar to those presented here in *C. elegans* may play a similar role in other organisms.

## Author Contributions

J.M., I.H., W.R.S. and Y.Z. designed the experiments. J.M., I.H., H.L., T.W. and S.B. performed experiments and analysed data. J.M., I.H., and W.R.S. wrote the manuscript and Y.Z., H.L., T.W., and S.B. read and critically revised the manuscript.

### Acknowledgements

We would like to acknowledge the Centre for Cellular Imaging at the University of Sweden, and the National Microscopy Infrastructure, NMI (NMI01125), for providing imaging facilities. This work was supported by grants from the Medical Research Council (MC-A023-5PB91), the Wellcome Trust (WT103784MA), National Institute of Health (W.R.S.), and the Swedish Research council (J.M.). Research in Zhang laboratory is supported by National Institutes of Health (NS115484, DC009852).

## Methods

### Key Resources

See Reagents and resources table

## Resources Availability

### Lead Contact

Further information and requests for C. elegans strains and plasmids is to be sent to and will be fulfilled by the Lead Contact William R Schafer

### Data and Code availability

Python scripts for TEVC analysis can be found at on GitHub at hiris25/TEVC-analysis-scripts. Aggregated data used for analysing TEVC data are available upon request from the Lead Contact.

## Experimental Model and Subject Details

### C. elegans

Unless otherwise specified, worms were maintained at 20°C on nematode growth medium (NGM) plates seeded with bacterial *E. coli* (strain OP50). Transgenic lines were generated by injection of plasmid DNA into the gonad of day 1 adult hermaphrodites. Offspring with stable arrays were selected. Mutant strains generated by CRISPR were outcrossed at least three times, mutant strains obtained from million mutation project (Thompson et al., 2013) or by UV transgene integration were outcrossed at least six times, all to our laboratory stock of wild-type (N2). A complete list of strains and transgenes used in this study see STAR Table.

### Xenopus laevis oocytes

Defolliculated *Xenopus laevis* oocytes were obtained from EcoCyte Bioscience (Dortmund, Germany) and maintained in ND96 (in mM: 96 NaCl, 1 MgCl_2_, 5 HEPES, 1.8 CaCl_2_, 2 KCl) solution at 16° C for 3-5 days.

## Method Details

### Phylogenetic analysis of *C. elegans* LGC genes

A set of 171 LGC protein sequences were submitted to MAFFT multiple alignment server (Katoh et al., 2018) using the L-INS-i method for sensitive alignment. The resulting alignment in CLUSTAL format was refined with trimal (Capella-Gutiérrez et al., 2009) using the parameters -gt 0.5 -w 7 to select only those alignment columns where considering the average of +/- 7 positions, 50% of sequences were devoid of gaps. The trimmed alignment file was converted to PHYLIP format using an online server (http://sequenceconversion.bugaco.com/converter/biology/sequences/). The alignment in PHYLIP format was submitted to the PHYML-SMS web server (Guindon et al., 2010) which predicted the LG +G+F as the optimal model for building a phylogenetic tree. Finally, RAxML v.8 was used to build a tree (Stamatakis, 2014) using the PHYLIP format trimmed alignment with the following parameters -f a -m PROTGAMMAILGF -p 12345 -x 12345 -# 1000, which runs the program using fast bootstrapping with the LG +G+F model at 1000 bootstraps. Phylogenetic tree visualisation was built in FigTree (Rambaut, 2016), collapsing multiple isoforms of the same gene together, and coloured by subgroup.

### Molecular biology

*C. elegans* cDNA sequences were cloned from wild-type N2 worm cDNA generated by reverse transcription PCR using Q5 polymerase (NEB, MA, USA) from total worm RNA. Ion channel cDNA sequences for *Xenopus* oocyte expression were cloned into the KSM vector backbone containing *Xenopus* β-globin UTR regions and a T3 promoter. *C. elegans* gDNA sequences were cloned from wild-type N2 worm gDNA. Transgene expression was verified by GFP or mKate2 expression, either fused to the protein, driven on the same plasmid after an intercistronic splice site (SL2 site), or co-injected with punc-122. Promoter sequences consisted of gDNA sequence approximately 2-3kb upstream of the start site of the gene. Subcloning was performed using HiFi assembly (NEB, MA, USA), IVA (*in-vivo* assembly, García-Nafría et al., 2016) or the Multisite Gateway Three-Fragment cloning system (Thermo Fisher Scientific, CA, USA) into pDESTR4R3II. Site-directed mutagenesis was performed using the KLD enzyme mix (NEB, MA, USA) or using IVA. For full list of primers used, see STAR Table.

### CRISPR/CAS9-mediated gene manipulation

Genetic modifications including deletions and point mutations were made by following the Dokshin et al. protocol (Dokshin et al., 2018). gRNA and ssODN were ordered from Sigma (Merck group, Darmstadt, Germany), a list of sequences is provided in STAR Table. Endogenous tagging of *lgc-50* with GFP was carried out using the SapTrap protocol (Dickinson et al., 2015; Schwartz and Jorgensen, 2016) where GFP was added in the cytosolic M3/4 loop of the protein, sequences of plasmids can be found in STAR Table.

### RNA synthesis and microinjection

5’ capped cRNA was synthesised *in vitro* using the T3 mMessage mMachine transcription kit according to manufacturer’s protocol (Thermo Fischer Scientific, CA, USA). RNA was then purified using the GeneJET RNA purification kit (Thermo Fischer Scientific, CA, USA) prior to cRNA injection. Defolliculated *Xenopus* oocytes were placed individually into 96 well plates and injected with 50nL of 500ng/µL RNA using the Roboinject system (Multi Channel Systems GmbH, Reutlingen, Germany). When two constructs were injected the total RNA concentration remained 500ng/µL, with a 1:1 ratio of the components. Injected oocytes were incubated at 16° C in ND96 until the day of recording, typically between 3-5 days post injection.

### Two-Electrode Voltage Clamp (TEVC)

Two-electrode voltage clamp recordings were carried out using the Robocyte2 recording system or a manual TEVC set up, using an OC-725D amplifier (Multi Channel Systems, Reutlingen, Germany) and paired with a custom-made recording chamber and agar bridges from reference and bath electrodes. Glass electrodes were pulled on a P-1000 Micropipette Puller (Sutter, Ca, USA) with a resistance ranging from 0.7-2 MΩ, pipettes, containing Ag|AgCl wires, were backfilled with a 3 M KCl solution for manual recordings and 1.5M KCl and 1 M acetic acid for Robocyte2 recordings. Oocytes were clamped at -60mV unless stated otherwise. Continuous recordings were taken during application of agonists and antagonists at 500 Hz. Data was recorded using WinWCP or RoboCyte2 control software, manual data was filtered at 10 Hz.

Dose response curves were calculated from the peak current during a 10s agonist stimulation in ND96 solution, with a 60s ND96 wash in between doses. Data was gathered over at least two occasions, using different batches of oocytes. Normalised dose response data was fitted to a nonlinear curve using a four parameters variable slope and the EC_50_ and Hill slope was calculated. All further recordings were carried out with the agonist at its EC_50_ concentration unless stated otherwise. Ion selectivity was determined using a voltage ramp protocol of 20mV/s ranging from -80mV to +60mV in the presence or absence of the primary agonist in three different solutions: ND96, NMDG (Na^+^ free) and Na Gluconate (low Cl^-^) solutions. Normalised ramp curves were fitted to a linear regression line and the x intercept was compared between solutions to calculate an average E_rev_ from 4-5 oocytes. Antagonist dose response curves were calculated from the peak current during a 10s agonist + antagonist stimulation in ND96 solution, with the agonist concentration remaining constant. Repeated agonist stimulus protocols were carried out by measuring the peak current during a 10s agonist stimulation at three wash intervals, 10s, 30s and 60s. Kinetic measurements were calculated from a 60s agonist perfusion.

### TEVC data analysis and plotting

Peak current was calculated using different software depending on origin of the data, manual recordings were analysed with WinWCP and Robocyte2 collected data was analysed with Stimfit or Robocyte2+. In all cases the peak current was taken during the window of interest.

Dose response and antagonist response curves were generated using custom-built python scripts (STAR table), which combined data from multiple recordings and normalised data by calculating I/Imax for each oocyte. Normalised mean, SD and n numbers where then imported into GraphPad where data was plotted and EC_50_ or IC_50_ values were calculated by fitting to the Hill equation using either three or four parameter slopes to obtain the highest degree of fit.

Ion selectivity analysis was performed using a custom-built python script (STAR table). Data was first normalised by calculating I/Imax for each oocyte and subtracting baseline currents from agonist induced currents in each solution. Non-linear quadratic line fitting was performed and reversal potential (E_Rev_) was calculated from the x intercept for each oocyte in each solution. Reversal potential shift (ΔE_Rev_) between ND96 and NMDG (Na_+_ free) and ND96 and Na Gluconate (low Cl_-_) solution was calculated for each oocyte and the individual values or mean, SD and n for each construct imported in GraphPad for plotting and statistical analysis. Statistically significant differences in ΔE_Rev_ were calculated in GraphPad using a 2way-ANOVA with Tukey’s correction for multiple comparisons. A selected representative trace, normalised by I/Imax and baseline subtracted, for each construct was also exported from python into GraphPad for plotting.

Repeated stimuli protocols were analysed by calculating peak current for each agonist window in Stimfit and exporting data to GraphPad, where data was plotted and significance was tested using 2way ANOVA with Tukey’s multiple comparison correction.

### Confocal and Cell ID

Worms were mounted onto an 2% agarose pad and immobilised using 75 mM NaAzide in M9. Images were acquired using a Leica SP8, with a 63x objective, for further analysis a collapsed z stack image was generated in Fiji/Image J. For identification of neurons carrying transgene expression different marker lines were used, as well as the multicolour reference worm NeuroPAL (Yemini et al., 2019) (for full list of lines see STAR Table).

### Image analysis

Confocal images of worms exposed to OP50 or the pathogenic bacteria *Serratia Marcescens* for 6h were analysed for calculating an intensity ratio between nerve ring and a posterior region next to the nerve ring using Fiji/Image J. Images were collapsed into one combined z-stack prior analysis. Intensity histograms across the entire worm was also analysed using the histogram tool in Fiji/Image J.

### Immunoprecipitation from *Xenopus* oocytes

IP experiments were performed on lysis from *Xenopus* oocytes expressing GFP tagged wild type LGC-50, LGC-50 Δ363-379 or uninjected oocytes as an IP control. Lysis buffer (0.3 M sucrose, 10 mM sodium phosphate (pH 7.4)), was supplemented with Halt Protease and phosphatase inhibitor 1:100 (Thermo Fischer Scientific) and NP40 at an end concentration of 0.5 % (Sigma Aldrich) immediately before usage. Oocytes were homogenised with 20 strokes in a glass homogeniser on day 3 after injection, keeping everything on ice. The lipid fraction was removed by centrifugation at 3000 x g at 4°C for 10 minutes. The lipid layer was removed and the remaining total lysate used for IP. 25 ml of equilibrated GFP-Trap MA beads (ChromoTek, GmbH) was incubated with 100 ml lysate at 4°C for 2h and then washed three times in TBS. Purified complexes were eluted from beads using Bolt LDS sample buffer and Bolt reducing agent (Thermo Fischer Scientific) at 70°C for 10 min before fractionated by SDS-PAGE (TGX Stain-free gel 4-20%, Bio-Rad).

### Immunoblotting

After SDS-PAGE electrophoresis the TGX stain-free gel was activated by UV in Gel Doc EZ (Bio-Rad) for quality control and the proteins transferred to a nitrocellulose membrane (Amersham Protran, 0.45 mm, ThermoFisher Scientific) with a wet transfer, (300 mA, 1 hour, Bio-rad). Membranes were blocked for 30 min in 5% milk and then incubated with primary antibody at 4°C overnight (anti-GFP HRP conjugated, A10260, diluted 1:1000 in TBS-T, ThermoFischer Scientific). The excessive unbound antibody was washed off the membrane using TBS-T before detection with Pierce ECL western blotting substrate (ThermoFischer Scientific). Blots were imaged using a ChemiDoc MP and Image Lab (Bio-Rad).

### Aversive olfactory training and learning assays

The aversive olfactory training with the pathogenic bacteria strain *P. aeruginosa* PA14 and the analysis of learning were performed similarly as previously described (Jin et al., 2016; Liu et al., 2018; Zhang et al., 2005). Adult *C. elegans* hermaphrodites cultivated under standard conditions were transferred onto a training plate, which was prepared by inoculating a NGM plate with fresh overnight culture of PA14 in NGM medium and incubating at 26°C for two days, or onto a control plate, which was prepared by inoculating a NGM plate with fresh overnight culture of OP50 in LB followed by two-day incubation at 26°C. The worms were trained for 4-6 hours at room temperature before learning assay.

To measure olfactory steering, a drop of 10 µL supernatant of a fresh overnight culture of PA14 or OP50 was put in the centre of a 10 cm NGM plate. One naive or trained worm was placed 1.5 cm away from the supernatant immediately before the recording started. The olfactory steering of the worm was recorded by a Grasshopper3-GS3-U3-120S6M-C camera (FLIR Integrated Imaging Solutions) at 7 frames per second and analysed by Wormlab (MBF Biosciences) and a MATLAB code (Liu et al., 2018). To measure the olfactory preference between PA14 and OP50, a two-choice preference assay was performed similarly as described (Jin et al., 2016; Liu et al., 2018). 20 µL of fresh overnight culture of PA14 and 20 µL of fresh overnight culture of OP50 were placed on the opposite sides of a standard chemotaxis plate (Bargmann et al., 1993) and dried on bench to form thin lawns before use. The concentration of the cultures was adjusted to optical density 600 = 1. Naive or trained worms were washed off respectively from the control or training plate and placed in the center of the testing plate, equidistance from the bacterial cultures, to test their olfactory preference between the two bacteria lawns. The number of the worms on each bacteria lawn was counted by the end of the assay to calculate learning index (Jin et al., 2016).

### Thermotaxis

The behaviour was briefly adapted from (Luo et al., 2014; Mori and Ohshima, 1995). In short, staged worms (L4 stage) were placed overnight at three different temperatures (15°C, 20°C and 25°C). The following day worms were tested on a specially built thermogradient equipment, that held a gradient from 15°C to 25°C vertically across the plate. The thermo stage had the size of a standard 96-well plate (128 ⨯ 85,5 mm), and rectangular four-well dishes (Nunc, ThermoFisher Scientific) filled with NGM agar were used for the testing. The worms were carefully washed in M9 before placed on testing plates to remove all bacteria. Worms were allowed to move freely over the temperature gradient for 1h after which the positions of the worms were scored.

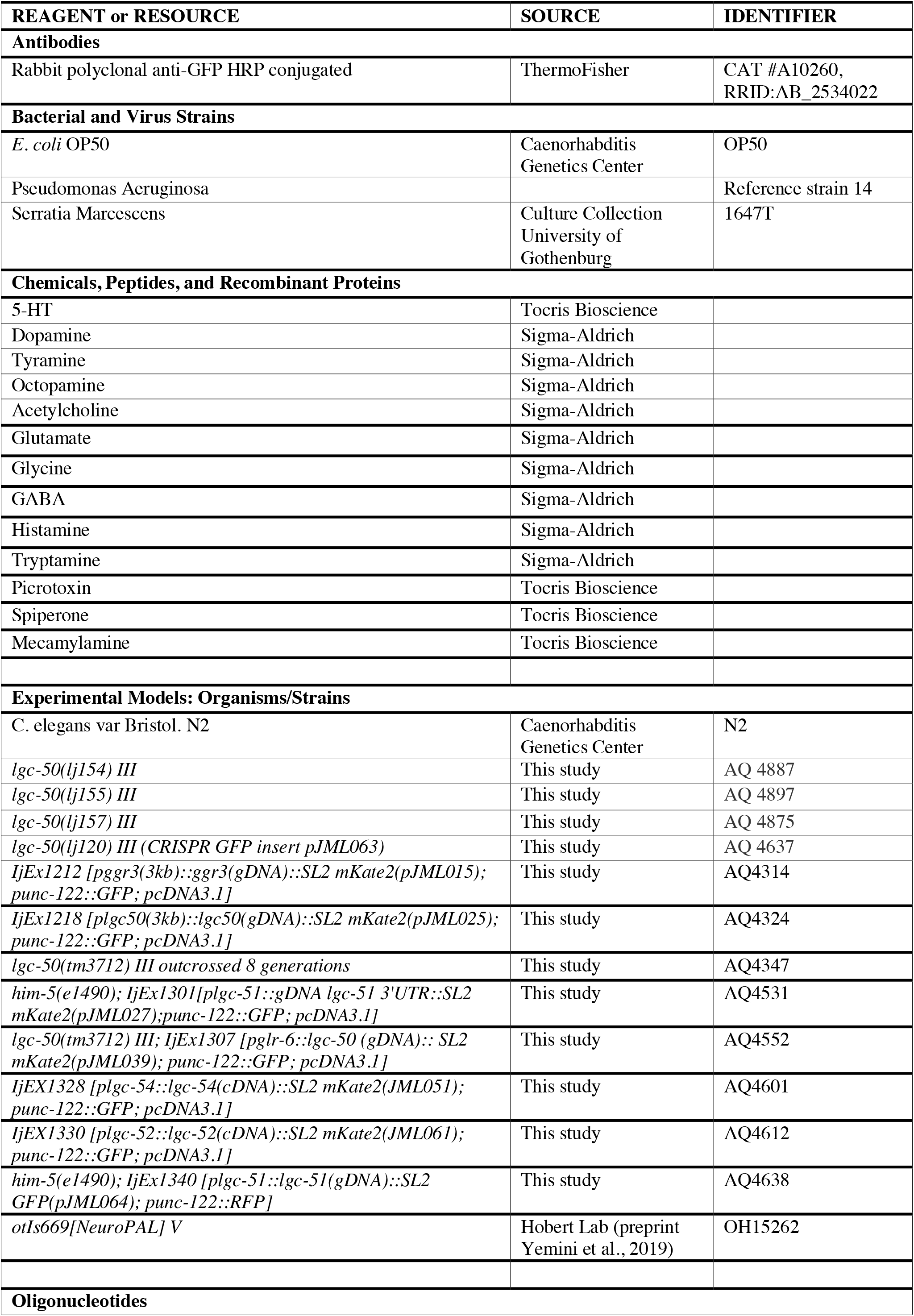
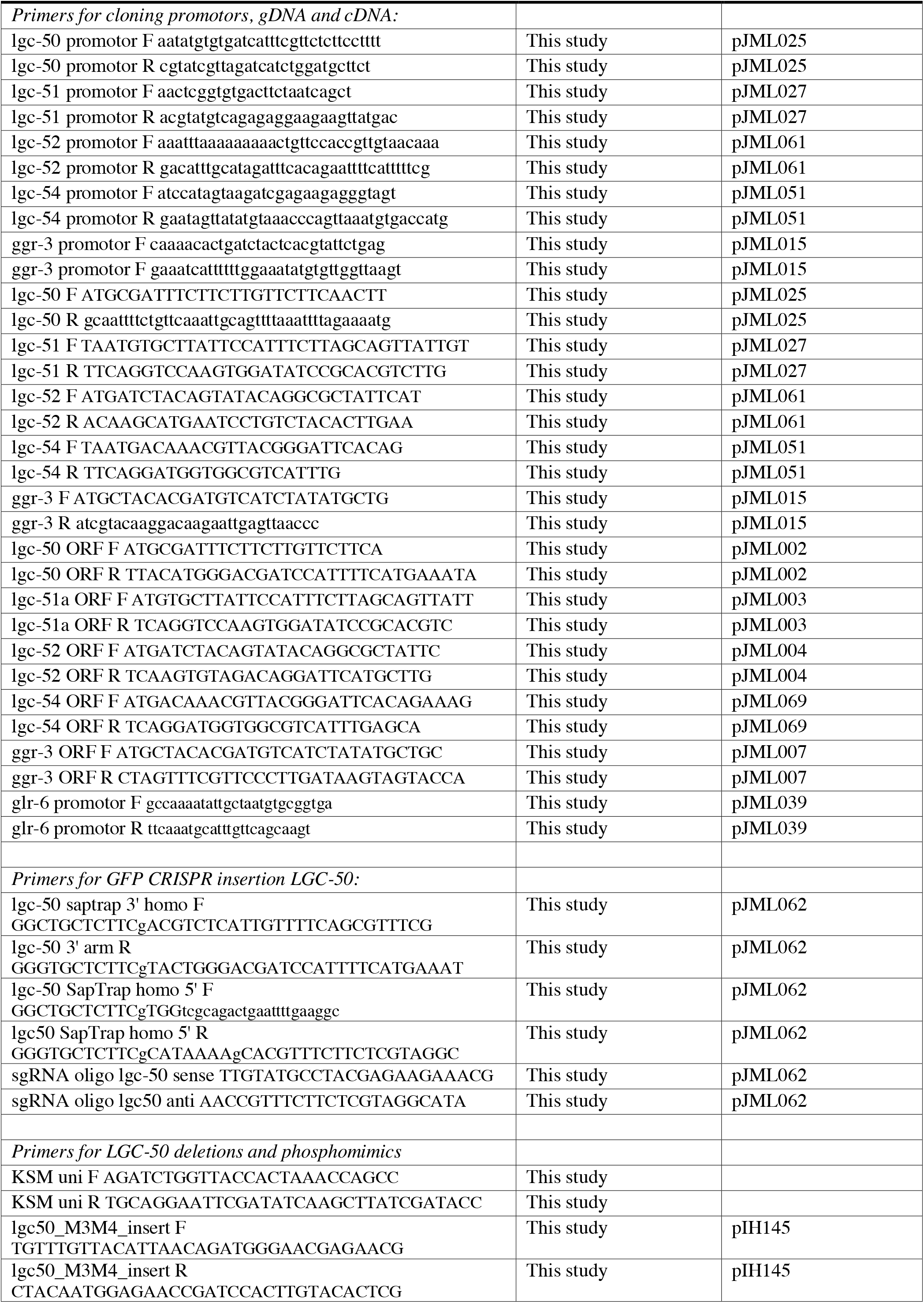
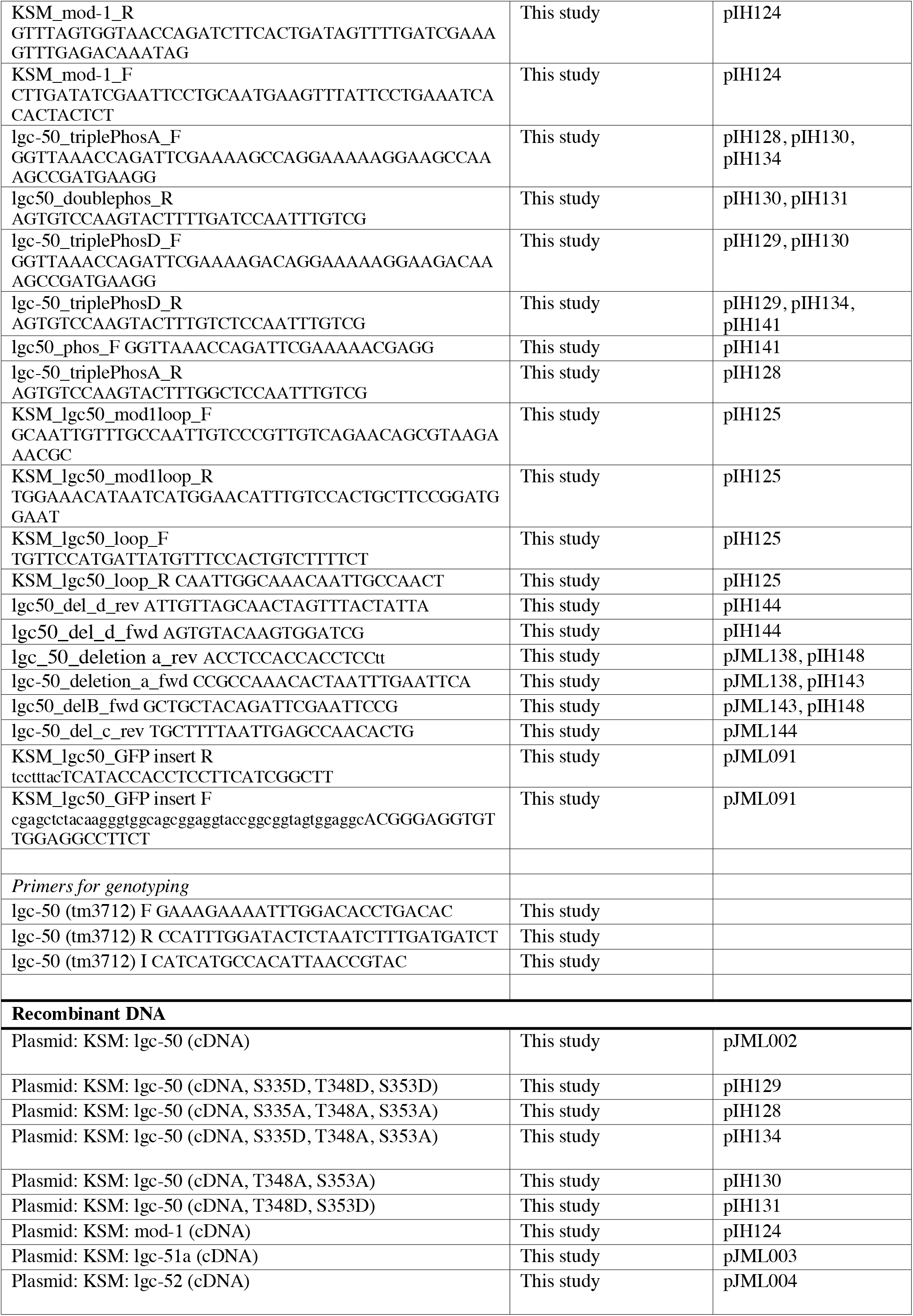
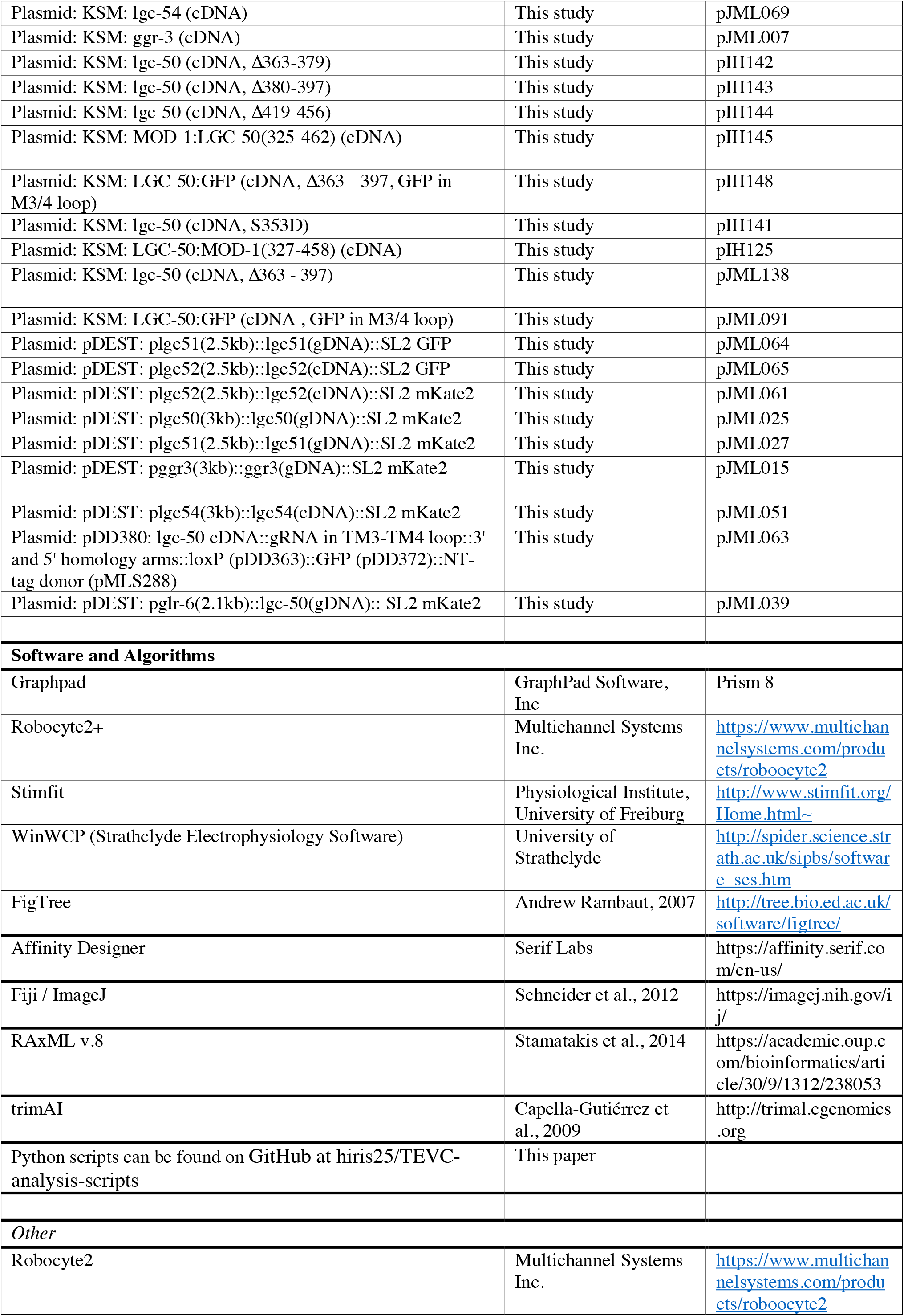
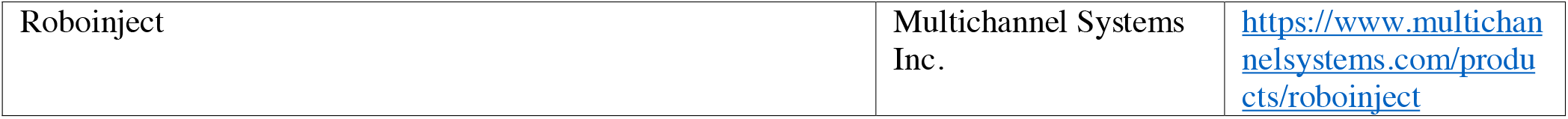

## Supplemental Information

*Table S1*

List of C. elegans genes used for phylogenetic analysis.

*Table S2*

Dataset generated using NetPhos 3.1 mapping potential phosphorylation sites in LGC-50.

**Figure S1.**
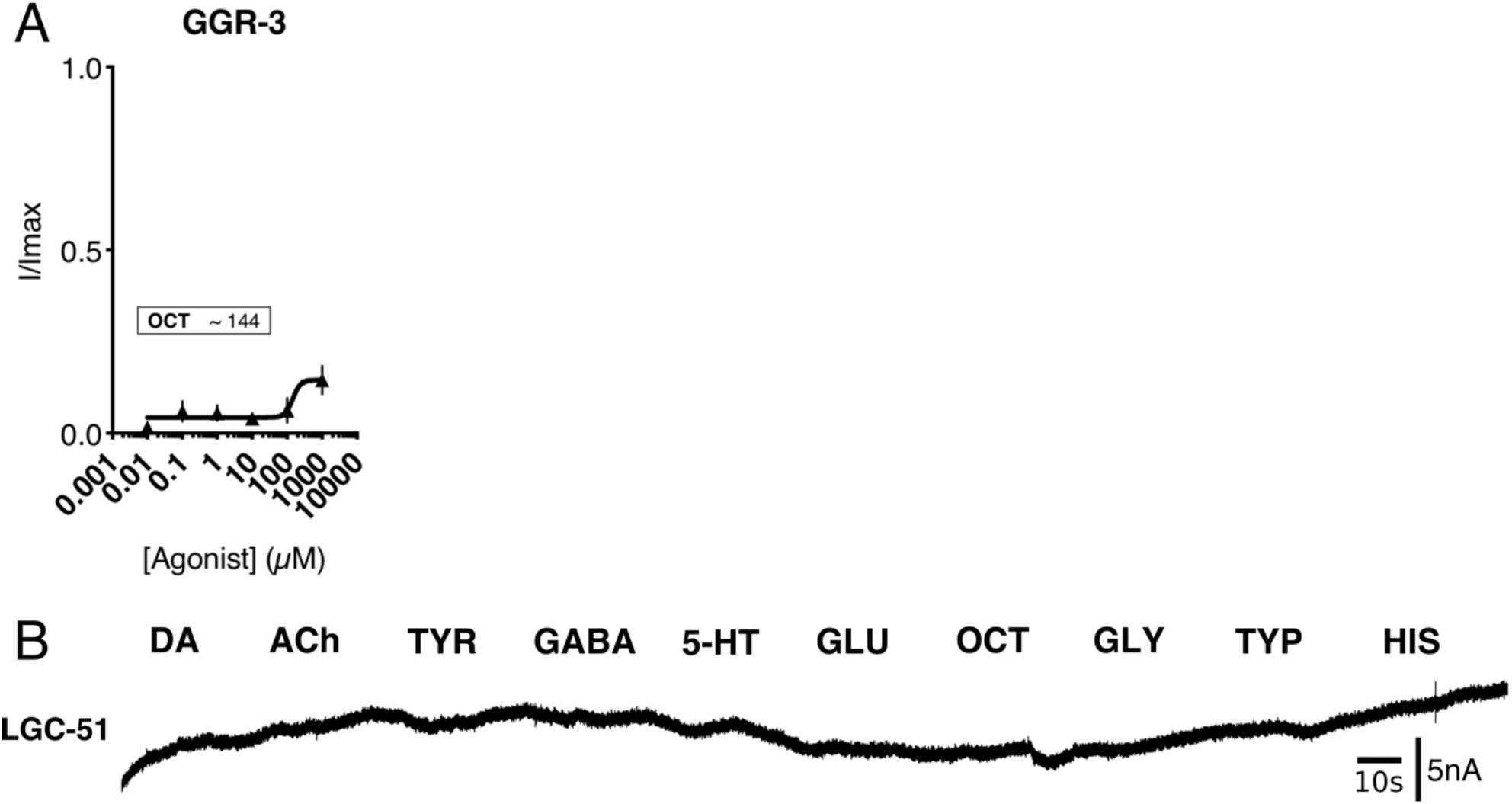
A. Dose response for octopamine application on GGR-3 B. Agonist panel application on LGC-51 when expressed as a homomer shows no opening of the channel to the possible agonists evaluated.

**Figure S2.**
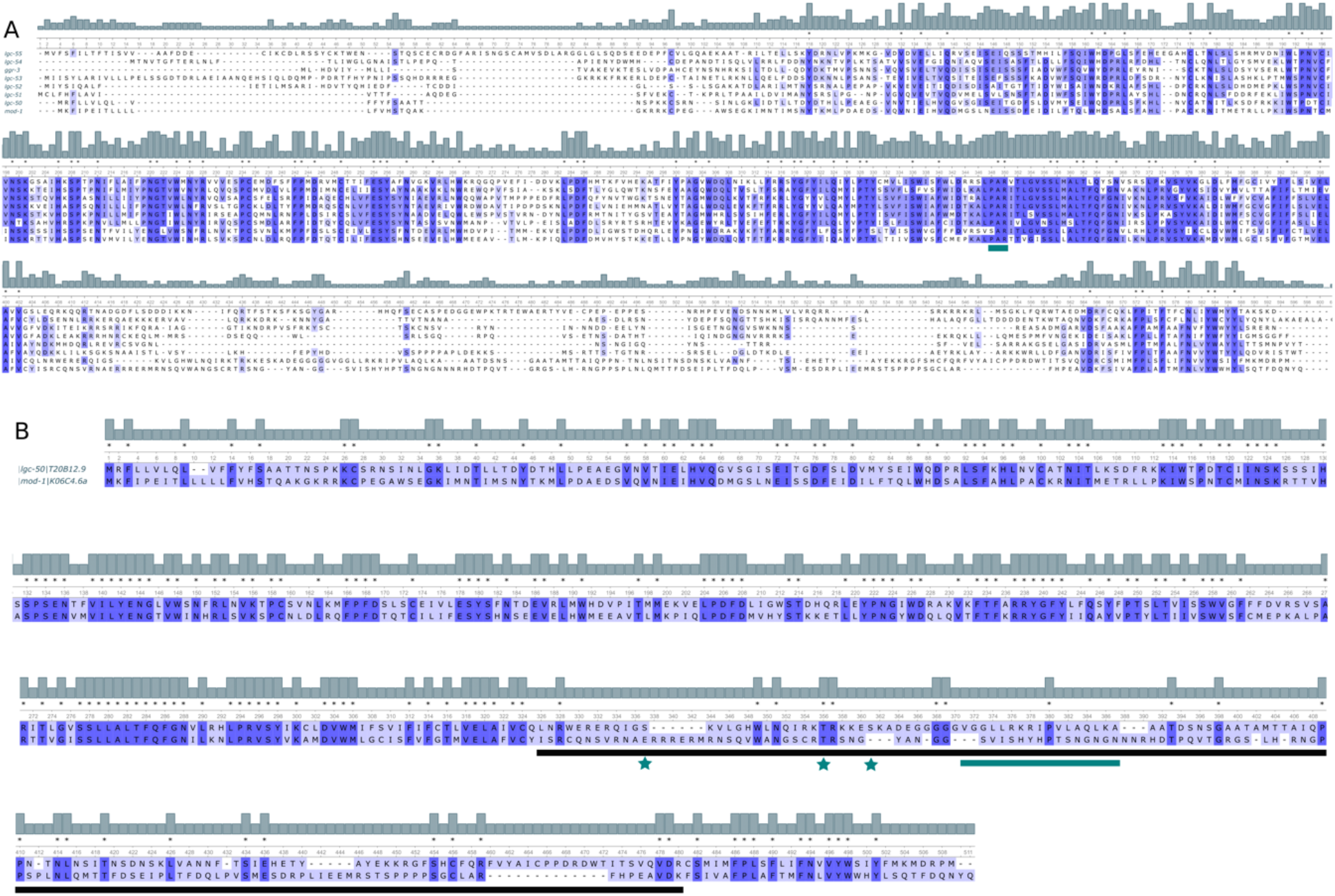
A. Alignment of aminergic LGCs shows at the green line conservation of the PAR motif (position 348) for all channels but LGC-50 that has a point mutation replacing proline with serine. B. Alignment of LGC-50 and MOD-1 shows high homology until the start of the M3/4 loop, which is indicated by the black line. The green stars indicate the predicted phosphorylation sites in LGC-50 (Table S2) and the green line marks the deleted region in LGC-50 (Δ363-379), that when deleted allows LGC-50 to traffic to the plasma membrane.

**Figure S3.**
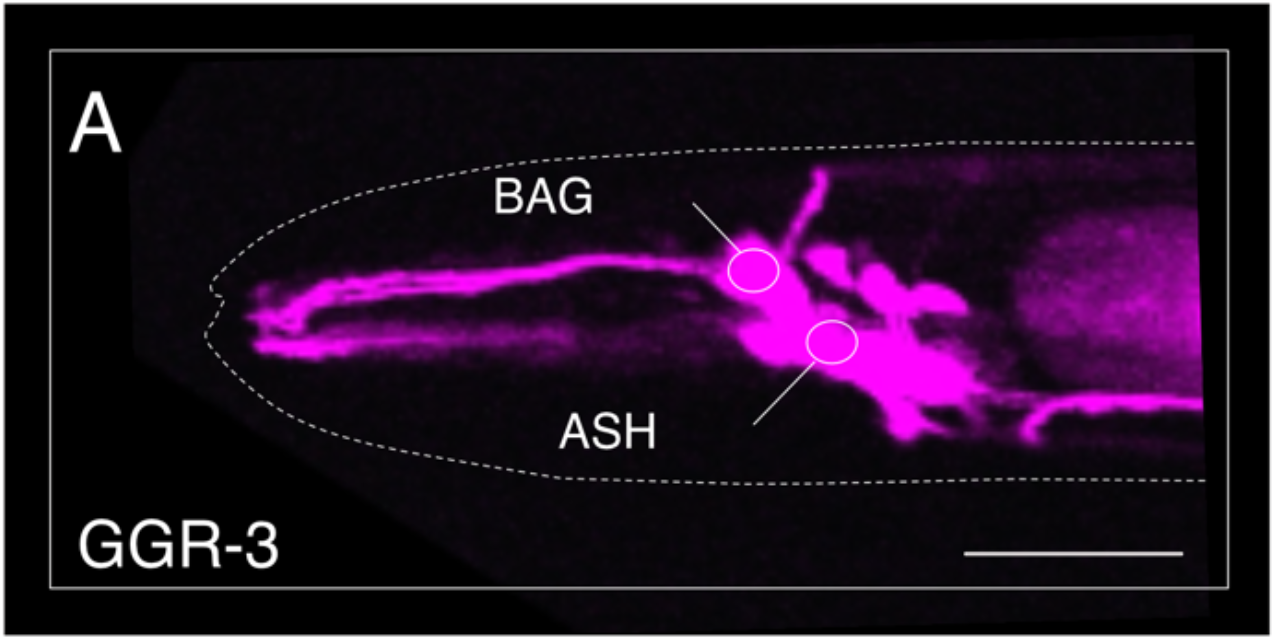
A. Confocal image of ggr-3:SL2 mKate denoting expression in the ASH and BAG neurons, as well as several yet unidentified neurons.

**Figure S4.**
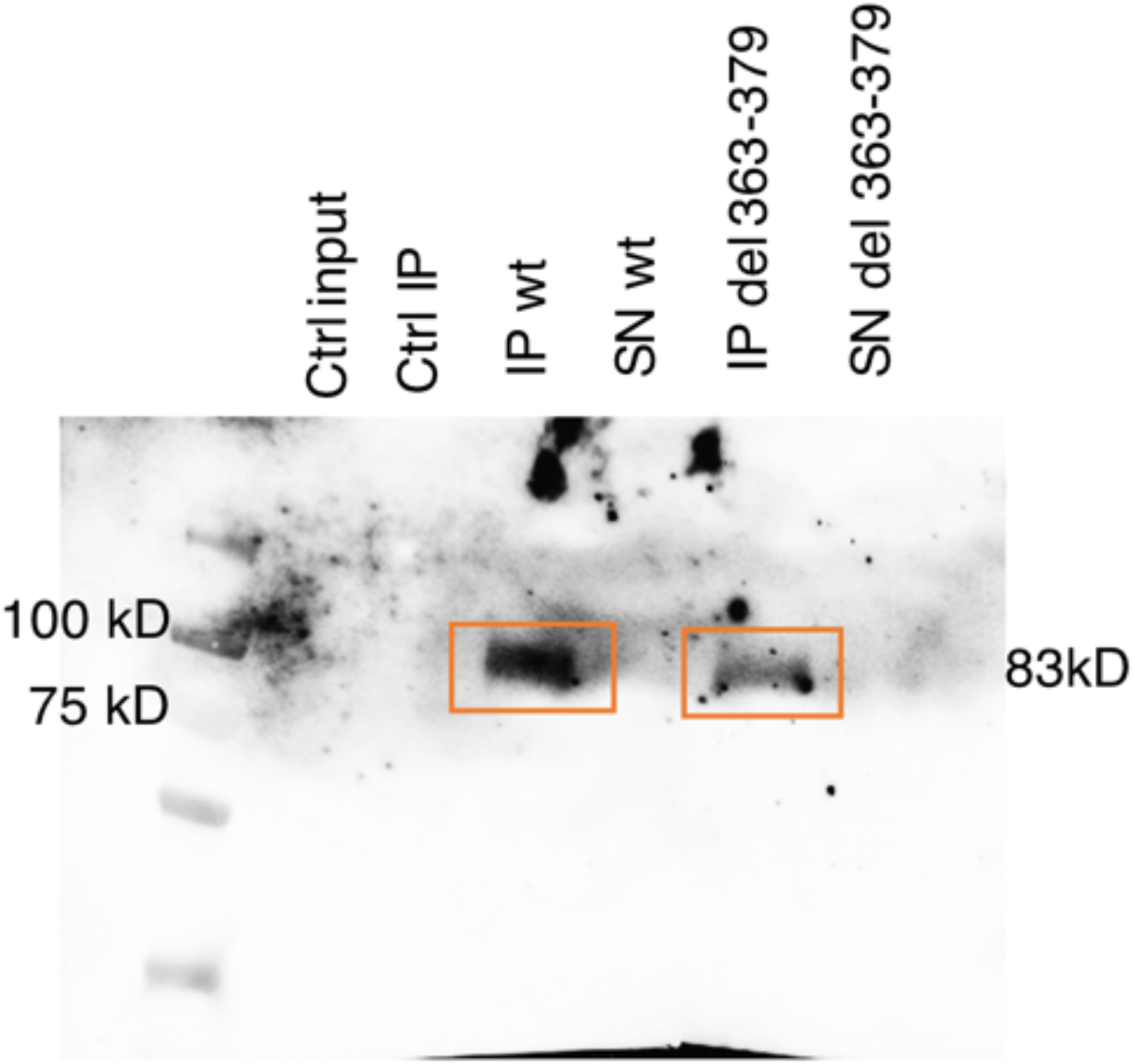
Western blot image after IP experiments in oocytes expressing wt LGC-50:GFP or LGC-50Δ363-379:GFP using anti-GFP antibody detection. Ctrl input: total lysate from untreated oocytes. IP: immunoprecipitation using anti-GFP magnetic beads, SN: supernatant after IP.

**Figure S5.**
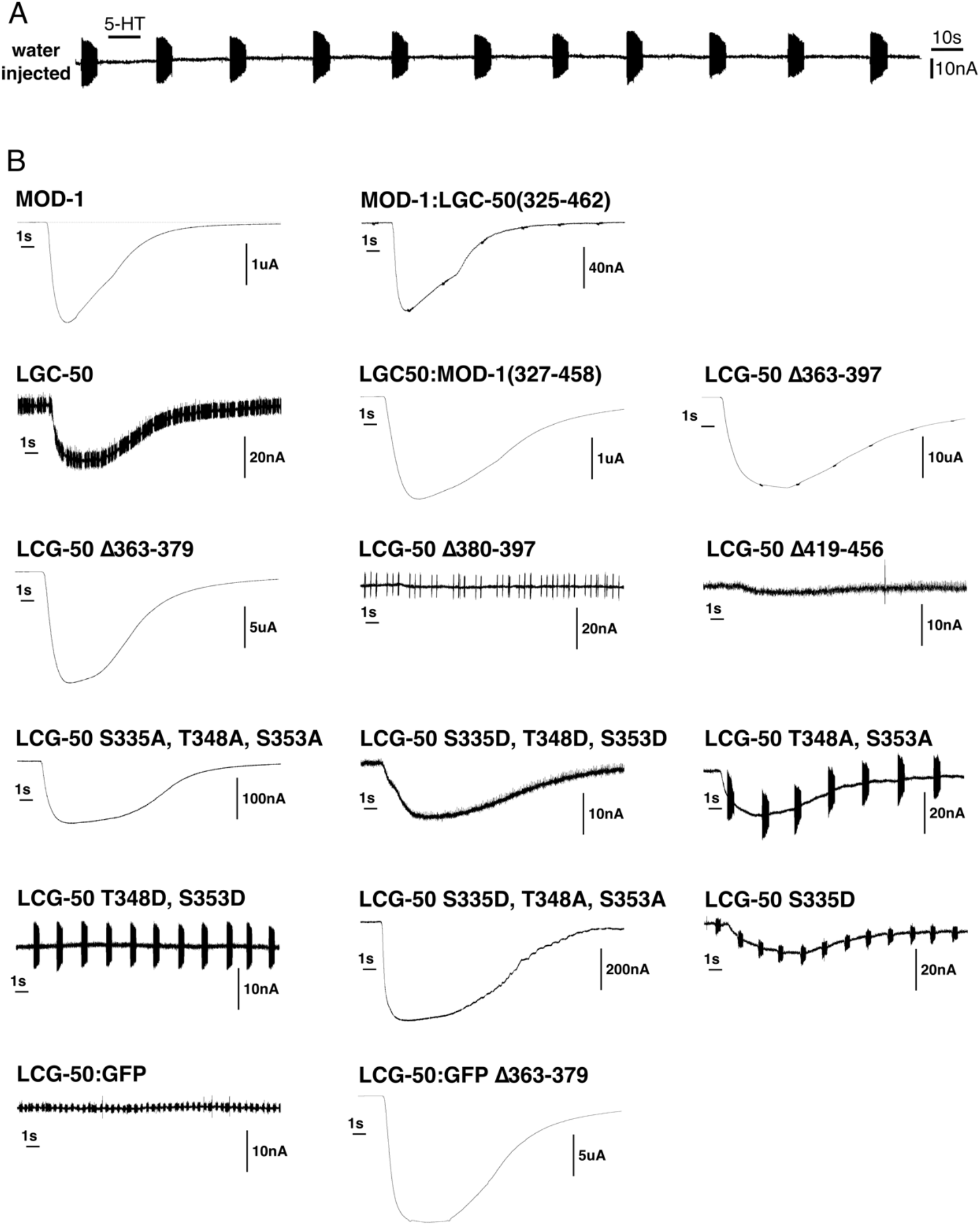
A. TEVC trace from water injected oocytes shows no response to 5-HT application. B. Representative traces from TEVC recordings in oocytes from different chimera proteins between MOD-1 and LGC-50, as well as several LGC-50 deletion and phosphomimic mutants.

**Figure S6.**
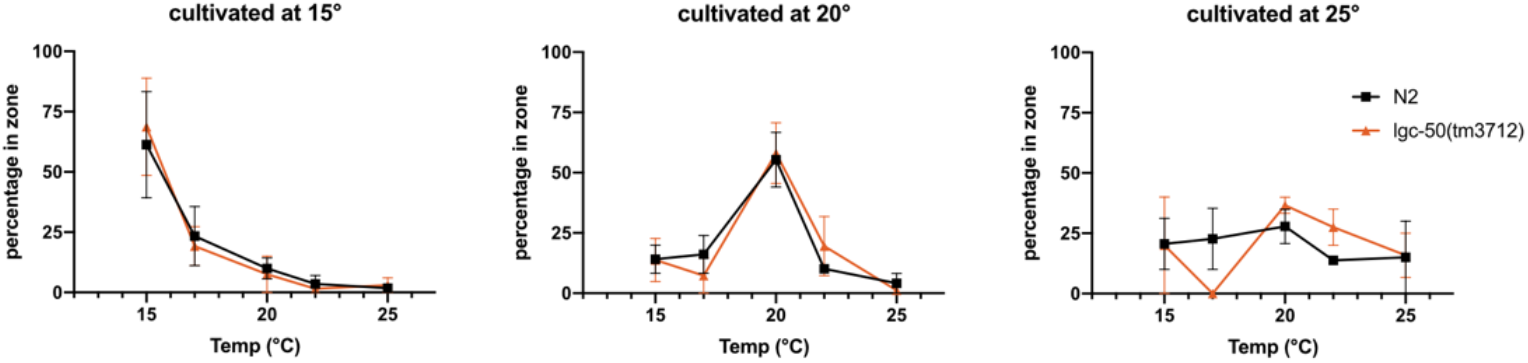
Thermotaxis data for comparing N2 and *lgc-50* mutant worms after cultivating the worms at three different temperatures overnight and performing 1h of thermotaxis testing the following day. No difference in thermotaxis behaviour was detected.

